# Mechanistic insights into structure-based design of a Lyme disease subunit vaccine

**DOI:** 10.1101/2024.10.23.619738

**Authors:** Kalvis Brangulis, Jill Malfetano, Ashley L. Marcinkiewicz, Alan Wang, Yi-Lin Chen, Jungsoon Lee, Zhuyun Liu, Xiuli Yang, Ulrich Strych, Maria-Elena Bottazzi, Utpal Pal, Ching-Lin Hsieh, Wen-Hsiang Chen, Yi-Pin Lin

## Abstract

The quality of protective immunity plays a critical role in modulating vaccine efficacy, with native antigens often not able to trigger sufficiently strong immune responses for pathogen killing. This warrants creation of structure-based vaccine design, leveraging high-resolution antigen structures for mutagenesis to improve protein stability and efficient immunization strategies. Here, we investigated the mechanisms underlying structure-based vaccine design using CspZ-YA, a vaccine antigen from *Borrelia burgdorferi*, the bacteria causing Lyme disease (LD), the most common vector-borne disease in the Northern Hemisphere. Compared to wild-type CspZ-YA, we found CspZ-YA_I183Y_ and CspZ-YA_C187S_ required lower immunization frequency to protect mice from LD-associated manifestations and bacterial colonization. We observed indistinguishable human and mouse antigenicity between wild-type and mutant CspZ-YA proteins after native infection or active immunization. This supports our newly generated, high-resolution structures of CspZ-YA_I183Y_ and CspZ-YA_C187S_, showing no altered surface epitopes after mutagenesis. However, CspZ-YA_I183Y_ and CspZ-YA_C187S_ favored the interactions between helices H and I, consistent with their elevated thermostability. Such findings are further strengthened by increasing ability of protective CspZ-YA monoclonal antibodies in binding to CspZ-YA at a physiological temperature (37°C). Overall, this study demonstrated enhanced intramolecular interactions improved long-term stability of antigens while maintaining protective epitopes, providing a mechanism for structure-based vaccine design. These findings can ultimately be extended to other vaccine antigens against newly emerging pathogens for the improvement of protective immunity.

## INTRODUCTION

Active immunization aims to trigger host immunity to eliminate pathogens during acute and subsequent infections [1]. Vaccination needs to be safe, and number of necessary immunizations should be limited, while still providing protection [2, 3]. However, some surface antigens, even in their recombinant forms, from infectious agents are less immunogenic, resulting in inefficient pathogen elimination [4]. That challenge sparks off multiple antigen engineering strategies to enhance antigenicity [5], one of which is the structure-based vaccine design [6, 7]. This strategy examines the structures of wild-type and mutant proteins seeking to improve protein stability and immunogenicity [6, 7]. Structure-based vaccine design has been demonstrated its suitability for antigen engineering against numerous pathogens to prevent infectious diseases, including the most recent outbreak of COVID-19 [8–10] (for review paper [11]). Nonetheless, the molecular mechanisms underlying the association between antigen stability and robust immunogenicity are still under investigation.

Lyme disease, also known as Lyme borreliosis, is the most common vector-borne disease in many parts of the Northern Hemisphere, with the number of human cases continuously rising (approximately 476,000 cases in the U.S. reported in 2022), with no effective prevention, such as human vaccines that are commercially available [12, 13]. As causative agents, multiple species of the spirochete bacteria, *Borrelia burgdorferi* sensu lato (also known as *Borreliella burgdorferi* or Lyme borreliae) are carried by infected *Ixodes* ticks and migrate to vertebrate hosts through tick bites [14]. Amongst those Lyme borreliae species, *B. burgdorferi* sensu stricto (hereafter *B. burgdorferi*) is the most prevalent human infectious Lyme borreliae species in North America while other human infectious species (e.g., *B. afzelii*, *B. garinii*, and *B. bavariensis*) are prevalent in Eurasia [15]. Upon introduction to hosts, Lyme borreliae colonize the tick bite sites in the skin and then disseminate through the bloodstream to distal organs, causing arthritis, carditis and/or neurological symptoms (i.e., neuroborreliosis) [16]. A human Lyme disease vaccine (LYMErix) was commercialized 20 years ago but then withdrawn from the market (detailed in [17, 18]). A second-generation vaccine is in clinical trials [19–22]. However, these vaccines target a Lyme borreliae protein, OspA, that is solely produced when bacteria are in the ticks but not in humans [23], thus preventing the development of any significant memory immune response against OspA [18]. Therefore, constant boosters of OspA-targeting vaccines is required to maintain protective levels of antibodies, challenging Lyme disease vaccine development [18].

Lyme borreliae produce other outer surface proteins, including CspZ (also known as BbCRASP-2 [24, 25]). CspZ promotes bacterial dissemination to distal tissues by evading the complement system, the first-line innate immune defense in vertebrate animals in the blood, through binding and recruiting a host complement inhibitor, factor H (FH) [24]. Although CspZ is not found in every Lyme borreliae strains [26], serologically confirmed and/or symptomatic human Lyme disease patients in North America and Eurasia all develop elevated levels of antibodies that recognize CspZ [27, 28]. These findings suggest the production of CspZ in most human infectious Lyme borreliae strains or species. Additionally, CspZ is only produced after Lyme borreliae invade vertebrate hosts, likely by triggering enhanced memory immune responses, underscoring the potential of employing this protein as a superior Lyme disease vaccine candidate [29, 30]. However, in mice, vaccination with the wild-type CspZ protein formulated with Freund’s adjuvant or aluminum hydroxide did not protect mice from Lyme borreliae colonization and Lyme disease-associated manifestations. One possibility is that CspZ’s protective epitopes are saturated by FH, which would not allow this protein to induce sufficient bactericidal antibodies to efficiently eliminate bacteria *in vivo*. We thus generated a CspZ-Y207A/Y211A mutant (CspZ-YA) that was shown to be selectively deficient in FH-binding [31], and such mutations thus lead to the exposure of the epitopes on this protein’s FH-binding sites[32, 33]. We demonstrated the protectivity of TiterMax Gold-adjuvanted CspZ-YA against tickborne infection of multiple human-infectious Lyme borreliae strains and species and correlated this with CspZ-YA-induced antibodies that uniquely recognize the epitopes surrounding the FH-binding site [33, 34]. These results and the availability of the high-resolution structure allows CspZ-YA as a model to test the concepts and mechanisms of action underlying structure-based vaccine design.

In this study, we engineered CspZ-YA by mutating the amino acids predicted through the structure-based vaccine design to test the efficacy of these CspZ-YA mutants in inducing bacterial killing and preventing Lyme disease-associated manifestations. We then examined the high-resolution structures and long-term stability of these CspZ-YA mutant proteins at 37 °C to identify the possible mechanisms underlying efficacy enhancement, advancing the understanding of the molecular basis for modern vaccine design strategies.

## RESULTS

### 1. Structure-based vaccine design identified amino acid residues enhancing the stability of CspZ-YA

To elucidate the structure basis of the effective CspZ-based vaccine antigens, CspZ-YA, we crystallized and obtained the structure of recombinant untagged version of this protein (1.90 Å) (**Fig. 1A**, **Table S1**). We then superimposed CspZ-YA with the previously determined crystal structures of the CspZ-FH complex [35]. The superimposed structures revealed that the Y211A mutation extended helix I by three residues (A211, K212, K213) (inlet figures in **Fig. 1A**). The extension of helix I and the Y207A mutation resulted in an altered conformation of the loop between helixes H and I (loop H/I, inlet figures in **Fig. 1A**). Such an orientation prevented R206 (located on the loop H/I) from interacting with E186 (inlet figures of **Fig. 1A**). Overall, the availability of this high-resolution structure built the foundation for the applications of further structure-based vaccine design of CspZ-YA [31].

**Figure 1.**
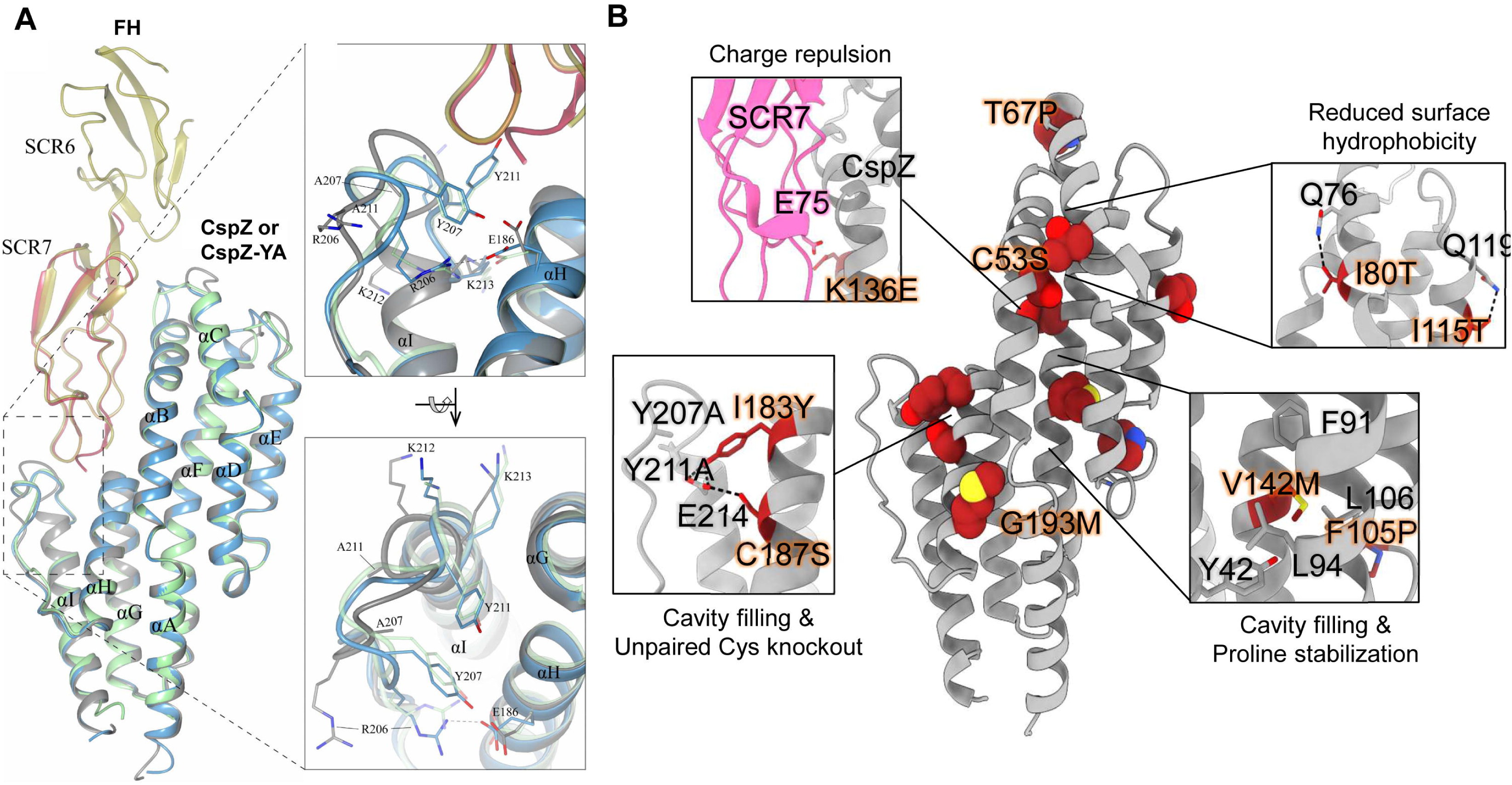
The high-resolution structure of CspZ-YA and the mutagenesis of amino acid residues in CspZ-YA by structure-based vaccine design. **(A)** The crystal structure of CspZ-YA (gray; PDB ID 9F1V; rmsd 1.9 Å) is superimposed with the structure of *B. burgdorferi* B31 CspZ (blue) from the CspZ/SCR6-7 (gold) complex (PDB ID 9F7I; rmsd 0.9 Å) and *B. burgdorferi* B31 CspZ (green) from the CspZ/SCR7 (red) complex (PDB ID 6ATG; rmsd 0.74 Å). The inlet figures show the loop region between helices H and I, highlighting residues Y207 and Y211 in CspZ from *B. burgdorferi* strain B31 and the mutated residues A207 and A211 in CspZ-YA. Residues K212 and K213 found in the loop region in CspZ and in the extended helix I in CspZ-YA are shown. The interaction between residues R206 and E186 in CspZ is further indicated. The structure is presented from top and side views. **(B)** Design landscape of CspZ-YA (PDB ID 9F1V) shown as a ribbon diagram with the side chains of the mutated amino acid residues shown as spheres. Insets highlight the position and side chains of selected stabilizing mutations. Side chains in each inset are shown as dark red sticks with sulfur atoms in yellow, nitrogen atoms in blue and oxygen atoms in red.

We then designed CspZ-YA variants to enhance the stability of CspZ-YA by substituting amino acid residues falling into one of four categories [6, 7]: (1) prolines at loop regions to decrease folding entropy (i.e., T67P, F105P), (2) polar residues to reduce surface hydrophobicity (i.e., I80T, I115T), (3) bulky hydrophobic residues to fill internal cavities, (i.e., V142M, I183Y, G193M), and (4) charge repulsions to disrupt FH binding (i.e., K136E) (**Fig. 1B**). The recombinant versions of CspZ-YA with T67P, I80T, F105P, I115T, K136E, V142M, I183Y, and G193M were produced in *E. coli* with histidine tags. Additionally, to avoid the formation of intermolecular disulfide bonds, resulting in the risk of protein aggregations [36–39], we substituted two cysteine residues, C53S and C187S to generate untagged CspZ-YA_C53S_ and CspZ-YA_C187S_ (**Fig. 1B**). The histidine-tagged and untagged proteins of CspZ-YA were also produced as control. [36–39]. We found that CspZ-YA_C53S_ was aggregated and insoluble (data not shown) while other CspZ-YA mutants were soluble and did not show differences in their secondary structures by circular dichroism, compared to CspZ-YA (**Fig. S1**). Therefore, all variants except CspZ-YA_C53S_ were moved forward to the following studies.

### 2. CspZ-YA_I183Y_ and CspZ-YA_C187S_ vaccinations triggered bactericidal antibodies and protect mice from Lyme disease infection with reduced immunization frequency

#### (i) Immunization with CspZ-YA_I183Y_ and CspZ-YA_C187S_ elicited robust borreliacidal antibody titers after two doses

To characterize the impact of these mutagenized amino acid residues on immunogenicity, we immunized mice with each of these CspZ-YA mutant proteins or CspZ-YA with different frequency (number of vaccination). The titers of anti-CspZ IgG in the sera at fourteen days post-last immunization (14 dpli) were determined (**Fig. 2A**). In any case, mice inoculated with any CspZ-YA proteins mounted significantly higher titers of CspZ IgG than those from PBS-inoculated control mice (**Fig. S2**). The IgG titers increased as the immunization frequency increased (**Fig. S2**). No significantly different titers were observed between the groups (**Fig. S2A to C**). These results indicate that the mutagenesis of these amino acid residues did not affect the overall IgG titers after vaccination. We then examined the ability of sera from 14 dpli to kill Lyme borreliae *in vitro* using *B. burgdorferi* strain B31-A3 as a model, the strain with the genotype (*ospC* type A) that is most prevalent in North America. When we compared these sera’s BA_50_ values, the dilution rate of the sera that kills 50% of bacteria, the BA_50_ values from any CspZ-YA proteins or variants increased when the immunization is more frequent (**Fig. 2B to G** and **Table S2**). The CspZ-YA mutants displayed no significantly different BA_50_ values from their parental CspZ-YA proteins after immunization once (**Fig. 2C**). Remarkably, CspZ-YA_I183Y_ and CspZ-YA_C187S_ triggered significantly (P<0.05) greater levels of BA_50_ values than the parental CspZ-YA proteins with twice or three times of the immunizations (**Fig. 2E and G**).

**Figure 2.**
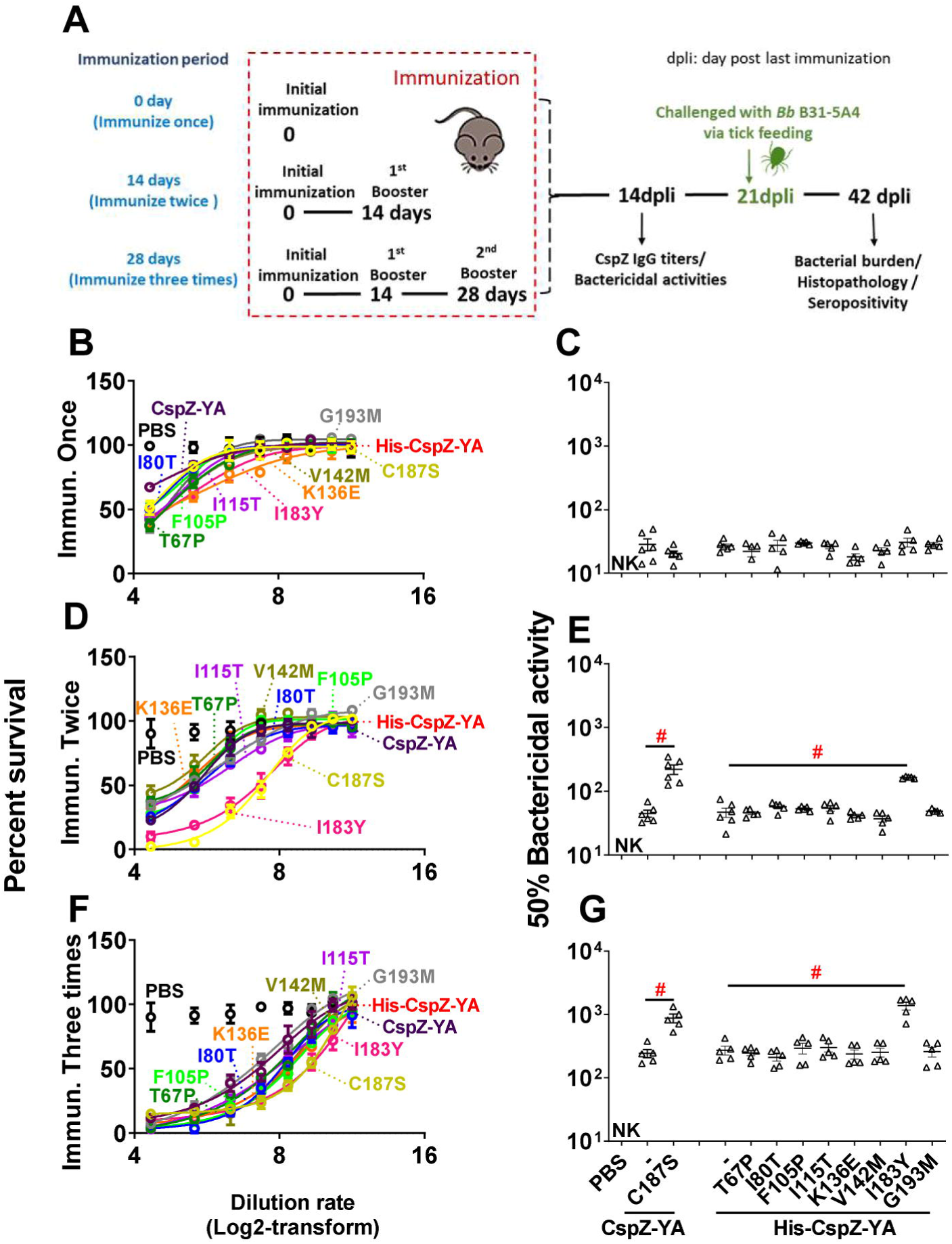
Mice immunized twice and three times with CspZ-YA_C187S_ or CspZ-YA_I183Y_ had sera with more robust levels of borreliacidal activity than CspZ-YA-vaccinated mice. **(A)** C3H/HeN mice received an inoculation with PBS (control) or immunization with CspZ-YA or the mutant proteins derived from this protein formulated with TitierMax Gold (TMG) at 0 day for the group of mice that were immunized once. The second group of mice that were immunized twice received the abovementioned proteins or PBS at 14 days after the initial immunization (dpii). The third group of mice that were immunized three times received the abovementioned proteins or PBS at 14 and 28 dpii. At 14 days post last immunization (14dpli), sera from these mice were collected for analyses of the titers of CspZ IgG and bactericidal activities. At 21 dpli, nymphal ticks carrying *B. burgdorferi* B31-A3 were placed on those mice and allowed to feed until repletion. Mice were sacrificed at 42 dpli for seropositivity, histopathology, and bacterial burden quantification. Mice inoculated with PBS and not fed on by nymphs were included as an uninfected control group. **(B to G)** Sera were collected at 14 dpli from C3H/HeN mice immunized **(B and C)** once, **(D and E)** twice, or **(F and G)** three times. These mice were immunized with PBS (control) or untagged CspZ-YA or its derived mutant proteins, or histidine tagged CspZ-YA (His-CspZ-YA), or its derived mutant proteins (Six mice for CspZ-YA- or CspZ-YA_C187S_-immunized mice whereas five mice for the rest of immunization groups of mice). These sera were serially diluted as indicated, and mixed with guinea pig complement and *B. burgdorferi* B31-A3 (5 × 10^5^ cells ml^-1^). After being incubated for 24 hours, surviving spirochetes were quantified from three fields of view for each sample using dark-field microscopy. The work was performed on three independent experiments. **(A, C, and E)** The survival percentage was derived from the proportion of serum-treated to untreated spirochetes. Data shown are the mean ± SEM of the survival percentage from three replicates in one representative experiment. **(B, D, and F)** The % borreliacidal dilution of each serum sample, representing the dilution rate that effectively killed 50% of spirochetes, was obtained from curve-fitting and extrapolation of Panel A, C, and E. Data shown are the geometric mean ± geometric standard deviation of the borreliacidal titers from three experiments. The exact values are shown in **Table S1.** PBS-inoculated mouse sera displayed no bactericidal activity (“NK”, no killing). Statistical significance (p < 0.05, Kruskal Wallis test with the two-stage step-up method of Benjamini, Krieger, and Yekutieli) of differences in borreliacidal titers between groups are indicated (“#”).

#### (ii) The I183Y and C187S mutations allowed CspZ-YA to protect mice from Lyme disease infection with fewer doses

We aimed to determine the ability of I183Y and C187S to reduce the protective immunization frequency of CspZ-YA against Lyme disease infection. We thus infected mice by permitting *Ixodes scapularis* nymphal ticks carrying *B. burgdorferi* B31-A3 to feed on the mice immunized with CspZ-YA or variant proteins under different immunization frequencies (**Fig. 2A**). We also included two controlled groups, PBS and lipidated OspA. At 42 dpli, we measured the bacterial burdens in different tissues and replete nymphs and detected the levels of IgG against C6 peptide, the commonly used Lyme disease serodiagnostic target (**Fig. 2A and Fig 3**) [40].

**Figure 3.**
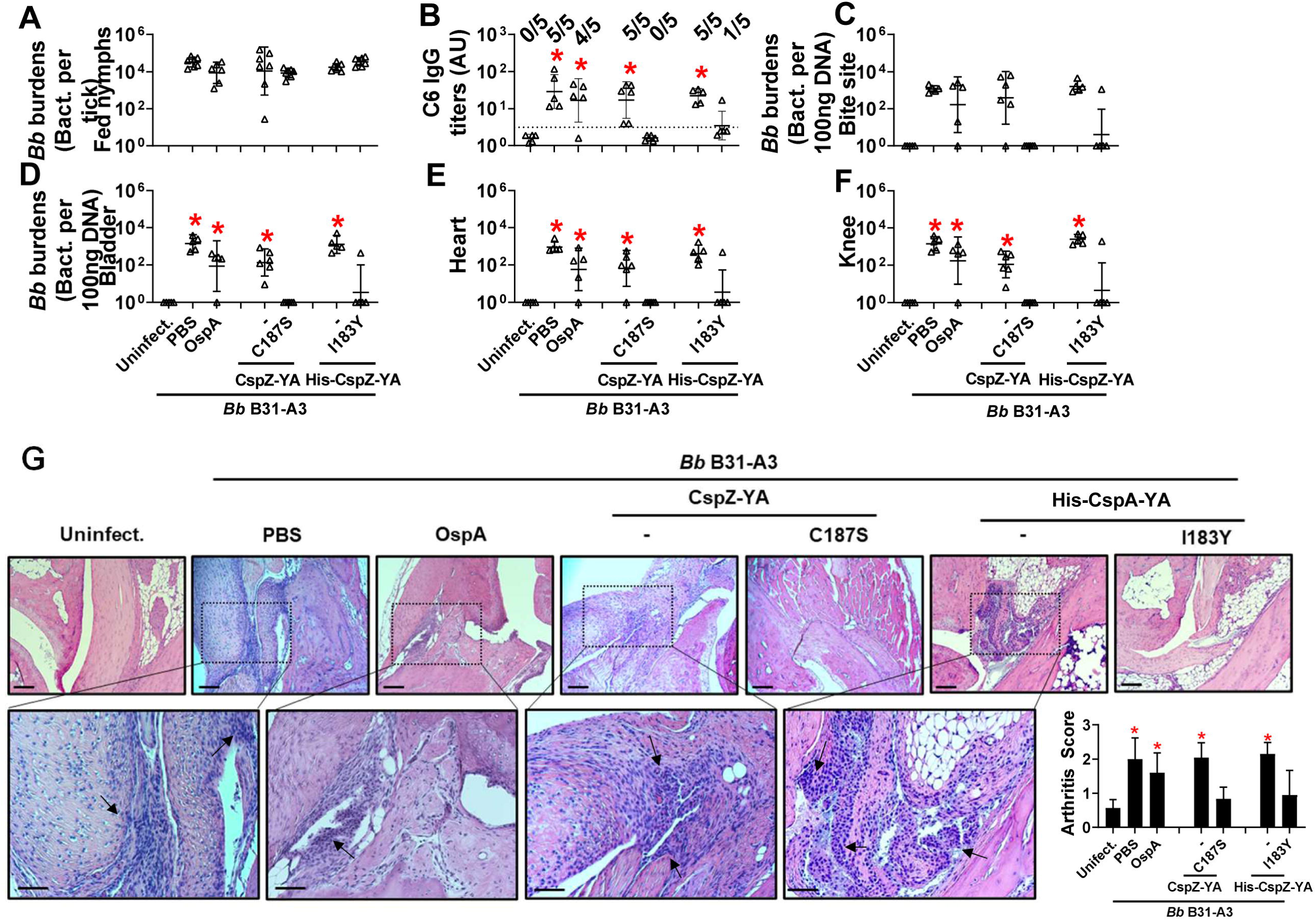
Immunizing twice with CspZ-YA_C187S_ or CspZ-YA_I183Y_ but not CspZ-YA protected mice from seroconversion, borrelial tissue colonization, and Lyme disease-associated arthritis. **(A to G)** Five PBS- or lipidated OspA (OspA)-, or histidine tagged CspZ-YA (His-CspZ-YA)- or CspZ-YA_I183Y_ (I183Y)-, or six untagged CspZ-YA- or CspZ-YAC187S (C187S)-immunized C3H/HeN mice that were immunized twice in the fashion described in Fig. 1 by indicated proteins. At 21days post last immunization, these mice were then fed on by nymphs carrying *B. burgdorferi* B31-A3. Mice inoculated with PBS that are not fed on by nymphs were included as an uninfected control group (uninfect.). **(B)** Seropositivity was determined by measuring the levels of IgG against C6 peptides in the sera of those mice at 42 days post last immunization using ELISA. The mouse was considered as seropositive if that mouse had IgG levels against C6 peptides greater than the threshold, the mean plus 1.5-fold standard deviation of the IgG levels against C6 peptides from the PBS-inoculated, uninfected mice (dotted line). The number of mice in each group with the anti-C6 IgG levels greater than the threshold (seropositive) is shown. Data shown are the geometric mean ± geometric standard deviation of the titers of anti-C6 IgG. Statistical significances (p < 0.05, Kruskal-Wallis test with the two-stage step-up method of Benjamini, Krieger, and Yekutieli) of differences in IgG titers relative to (*) uninfected mice are presented. (A, C to F) *B. burgdorferi* (*Bb*) burdens at **(A)** nymphs after when feeding to repletion or **(C**) the tick feeding site (“Bite Site”), **(D)** bladder, **(E)** heart, and **(F)** knees, were quantitatively measured at 42 days post last immunization, shown as the number of *Bb* per 100ng total DNA. Data shown are the geometric mean ± geometric standard deviation of the spirochete burdens from each group of mice. Asterisks indicate the statistical significance (p < 0.05, Kruskal Wallis test with the two-stage step-up method of Benjamini, Krieger, and Yekutieli) of differences in bacterial burdens relative to uninfected mice. **(G)** Tibiotarsus joints at 42 days post last immunization were collected to assess inflammation by staining these tissues using hematoxylin and eosin. Representative images from one mouse per group are shown. Top panels are lower-resolution images (joint, ×10 [bar, 160 µm]); bottom panels are higher-resolution images (joint, 2×20 [bar, 80 µm]) of selected areas (highlighted in top panels). Arrows indicate infiltration of immune cells. **(Inlet figure)** To quantitate inflammation of joint tissues, at least ten random sections of tibiotarsus joints from each mouse were scored on a scale of 0-3 for the severity of arthritis. Data shown are the mean inflammation score ± standard deviation of the arthritis scores from each group of mice. Asterisks indicate the statistical significance (p < 0.05, Kruskal Wallis test with the two-stage step-up method of Benjamini, Krieger, and Yekutieli) of differences in inflammation relative to uninfected mice.

We found that the fed nymphs from the mice inoculated with PBS, OspA, CspZ-YA, or CspZ-YA_C187S_ under any immunization frequency accounted for similar levels of bacterial burden (**Fig. 3A, S3A and S3G**). This is in agreement with prior findings that vaccination three times with OspA or CspZ-YA does not eliminate *B. burgdorferi* in fed ticks [33, 41]. Mice immunized once with any tested antigens were all seropositive (**Fig. S3B**) and had significantly greater bacterial loads in indicated tissues than uninfected mice (**Fig. S3C to F**). In contrast, mice immunized three times with any tested antigen and infected with *B. burgdorferi* were seronegative for C6 IgG (**Fig. S3H**) and showed no significantly different levels of bacterial burdens in tissues than uninfected mice (**Fig. S3I to L**). However, while all mice inoculated twice with PBS, OspA, CspZ-YA (untagged or histidine tagged), or were seropositive for C6 IgG, all CspZ-YA_C187S_-inoculated mice were seronegative (**Fig. 3B**). CspZ-YA_C187S_-inoculated mice showed no significantly different levels of bacterial burdens at tissues, compared to uninfected mice (**Fig. 3C to F**). We also included the mice immunized twice with CspZ-YA_I183Y_ and found that four out of five CspZ-YA_I183Y_-inoculated mice were seronegative (**Fig. 3B**) and had no significantly different levels of bacterial loads at tissues from those in uninfected mice (**Fig. 3C to F**). These results identified that two doses of CspZ-YA_C187S_ or CspZ-YA_I183Y_ but not CspZ-YA and OspA prevented colonization with *B. burgdorferi* and Lyme disease seroconversion. We further determined the severity of Lyme disease-associated arthritis in mice after two immunizations by histologically examining the mouse ankles. At 21 dpli, in OspA-, CspZ-YA-(untagged or histidine tagged), or PBS-inoculated mice, we found an elevated number of neutrophils and monocytes infiltrating the tendon, connective tissues, and muscles (arrows in **Fig. 3G**). However, similar to uninfected mice, there was no inflammatory cell infiltration in the joints from CspZ-YA_I183Y_- or CspZYA_C187S_-vaccinated mice (**Fig. 3G**). Overall, the mutagenesis of I183Y and C187S allows CspZ-YA vaccination to prevent Lyme disease infection with fewer immunizations.

### 3. Enhanced intramolecular interaction to stabilize protective epitopes of CspZ-YA_I183Y_ and CspZ-YA_C187S_ provides mechanisms underlying structure-based vaccine design

#### (i) The I183Y and C187S mutations did not alter the surface epitopes of CspZ-YA

One hypothesis to address the mechanisms underlying mutagenesis-mediated efficacy enhancement is that CspZ-YA_I183Y_ and CspZ-YA_C187S_ contain novel epitopes, distinct from CspZ-YA. We thus obtained the sera Lyme disease seropositive patients with elevated levels of CspZ IgGs (36 out of 38 serum samples have elevated levels of CspZ IgGs, **Fig. S4**). We then tested this hypothesis by comparing the ability of CspZ IgGs in these sera to recognize CspZ-YA, CspZ-YA_I183Y_, and CspZ-YA_C187S_. We found no significant difference between recognition of CspZ-YA, CspZ-YA_I183Y_, and CspZ-YA_C187S_ (**Fig. 4A**, two-tier pos.) whereas there was minimal detection with sera from seronegative humans from non-endemic areas (**Fig. 4A**, neg. ctrl.). A significantly positive correlation was detected for individual patient serum sample to recognize CspZ-YA, CspZ-YA_I183Y_, and CspZ-YA_C187S_ (**Fig. 4B to D**). We also compared the CspZ-IgGs produced in mice immunized twice with CspZ-YA (tagged or untagged), CspZ-YA_I183Y_, and CspZ-YA_C187S_ to recognize each of these proteins in the same fashion. We found that similar levels of recognition by CspZ-YA, CspZ-YA_I183Y_, and CspZ-YA_C187S_ for the sera from each immunization group of mice, and such levels of recognition are greater than those from PBS-inoculated control mice (**Fig. 4E**). Additionally, when combining the values of recognition from different immunization groups of mice, we observed a significantly positive correlation for those sera to recognize CspZ-YA, CspZ-YA_I183Y_, and CspZ-YA_C187S_ (**Fig. 4F to H**). Such indistinguishable human or mouse CspZ IgGs recognition by CspZ-YA, CspZ-YA_I183Y_, and CspZ-YA_C187S_ does not support the hypothesis that that mutagenesis of I183Y and C187S changes epitopes of CspZ-YA. We also crystallized and obtained the structure of the CspZ-YA_C187S_ (2.00 Å) and did the AlphaFold prediction for CspZ-YA_I183Y_ to examine the impact of those mutations for the epitopes *in silico*. Superimposed structures of CspZ-YA, CspZ-YA_C187S_, and CspZ-YA_I183Y_ showed no significant difference in surface epitopes (**Fig. 4I**). This is consistent with the fact that I183 and C187 are not surface-exposed but instead are buried between helices H and I of CspZ-YA (**Fig. 4J**). Taken together, our structural and immunogenicity results demonstrated no alteration of surface epitopes after the mutation of I183 and C187 in CspZ-YA.

**Figure 4.**
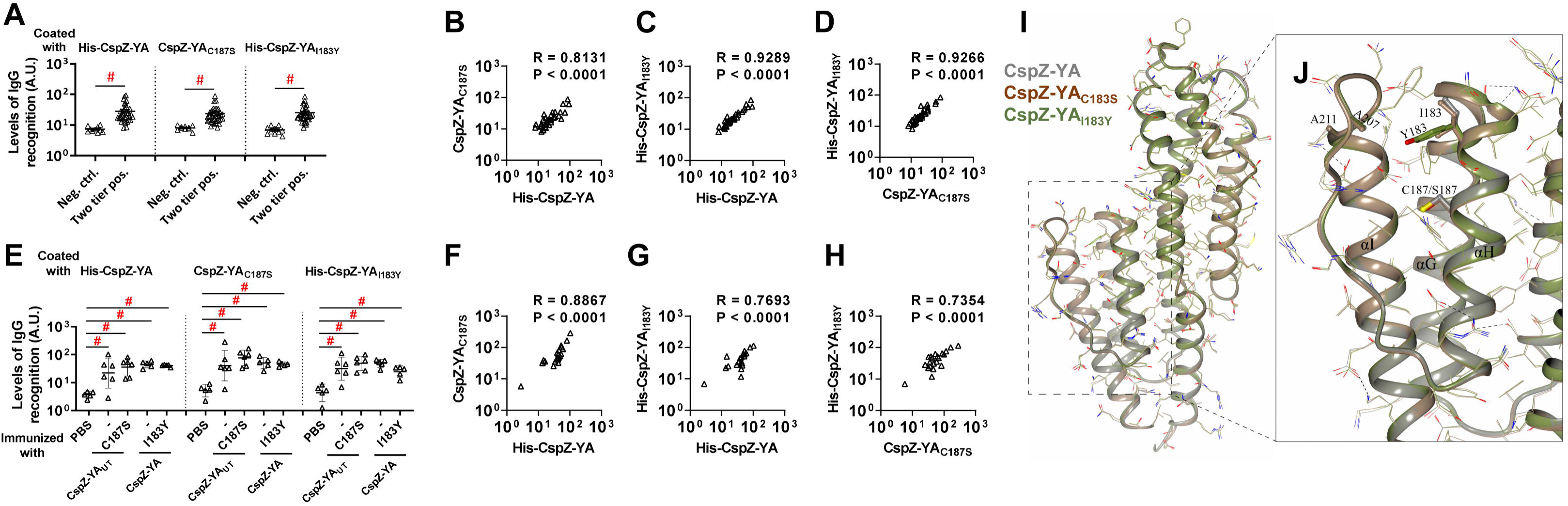
CspZ antibodies originating from humans or mice recognized CspZ-YA_C187S_ or CspZ-YA_I183Y_ at indistinguishable levels from CspZ-YA. **(A to D)** Sera from 36 patients with both seropositive for Lyme disease (“Two tier positive”; Positive in two tier test) in **Fig. S4** were included. **(A)** These sera were applied to determined their levels of recognition to histidine tagged CspZ-YA (His-CspZ-YA) or CspZ-YA_I183Y_ (His-CspZ-YA_I183Y_), or untagged CspZ-YA_C187S_ or using ELISA as described in the section “ELISA” in Materials and Methods. Ten serum samples from humans residing in non-endemic area of Lyme disease were included as negative control. Data shown are the geometric mean ± geometric standard deviation of levels of recognition in each group of serum samples. Statistical significance (p < 0.05, Kruskal Wallis test with the two-stage step-up method of Benjamini, Krieger, and Yekutieli) of differences in levels of recognition by groups are indicated (“#”).**(E to H)** Sera from five histidine tagged His-CspZ-YA or CspZ-YA_I183Y_ (I183Y)-, or six untagged CspZ-YA- or CspZ-YAC187S (C187S)-immunized C3H/HeN mice that were immunized twice in the fashion described in Fig. 1 were collected at 14dpli. PBS-inoculated mice were included as control. For each serum sample, the levels of its recognition by histidine tagged CspZ-YA, CspZ-YA_C187S_ or CspZ_I183Y_ were measured using ELISA. Data shown are the geometric mean ± geometric standard deviation of levels of recognition. Statistical significance (p < 0.05, Kruskal Wallis test with the two-stage step-up method of Benjamini, Krieger, and Yekutieli) of differences in levels of recognition by groups are indicated (“#”). For each serum sample originated from **(B to D)** humans or **(F to H)** mice, the values representing the levels of recognition by **(B and F)** CspZ-YA vs. CspZ-YAC187S, **(C and G)** CspZ-YA vs. CspZ-YAI183Y, or **(D and H)** CspZ-YAC187S vs. CspZ-YAI183Y were plotted. The correlation of these values derived from recognition by each of indicated CspZ-YA proteins was quantitatively determined using Spearman analysis and shown as R values. P values are also shown to demonstrate the statistical significance (p < 0.05, Spearman analysis) of the correlation between indicated values in X- and Y-axis in panel B to D and F to H. **(I)** Superimposed crystal structures of CspZ-YA (gray; PDB ID 9F1V), CspZ-YA_C187S_ (brown; PDB ID 9F21) and the predicted structure of CspZ-YA_I183Y_ (green). Side chains as thin bonds in all three proteins are illustrated. **(J)** Shown is the region in CspZ-YA, CspZ-YA_C187S_ and CspZ-YA_I183Y_ where mutations (A207, A211, C187, S187, I183, and Y183) were introduced. Residues associated with mutations are illustrated as thick bonds, but all other residues in all three proteins are represented as thin bonds. All the interactions observed between the amino acid side chains in CspZ-YA are indicated as dotted lines.

#### (ii) The I183Y and C187S mutations resulted in enhanced interactions between helix H and I of CspZ-YA proteins

Both I183 and C187 are located on and buried in helix H, raising the possibility that efficacy improvement can be attributed to the stabilization and/or enhancement of intramolecular interactions. We attempted to use protein crystal structures to investigate this possibility by comparing the electron density map surrounding C187 and S187 of CspZ-YA and CspZ-YA_C187S_, respectively. We found a water molecule present in the space between the helix I and C187 of CspZ-YA, as well as between helix I and S187 of CspZ-YA_C187S_ (see the arrow in **Fig. 5A and B**). While that water molecule does not impact the intramolecular interactions in CspZ-YA (**Fig. 5A**), that water molecule coordinates the interactions with S187 on the helix H and E214 on the helix I via hydrogen bonding in CspZ-YA_C187S_ (dotted lines in **Fig. 5B**). Additionally, the high-resolution structure of CspZ revealed a hydrophobic core formed among Y207, F210, and Y211 (**Fig. 5C**). Unlike CspZ, a cavity was found in CspZ-YA due to the replacement of two bulky and non-polar amino acid residues, Y207 and Y211, by alanine, as well as the orientation of F210 away from the hydrophobic core (the red highlight in **Fig. 5D**). Additionally, the altered conformation in CspZ-YA prevent**s** R206 from interacting with E186, exacerbating the cavity-mediated structural instability (**Fig. 5D**). Unlike CspZ-YA, we found that the AlphaFold predicted structure of CspZ-YA_I183Y_ showed that the cavity is filled by a bulky, non-polar residue (i.e., Y183), restoring the hydrophobic core between helix H and I (**Fig. 5E**). This is consistent with the intention of achieving a more stabilized structure of CspZ-YA_I183Y_ (**Fig. 1B**). Taken together, the structural evidence here suggests I183Y and C187S mutagenesis facilitates the helix H-I interactions of CspZ-YA proteins.

**Figure 5.**
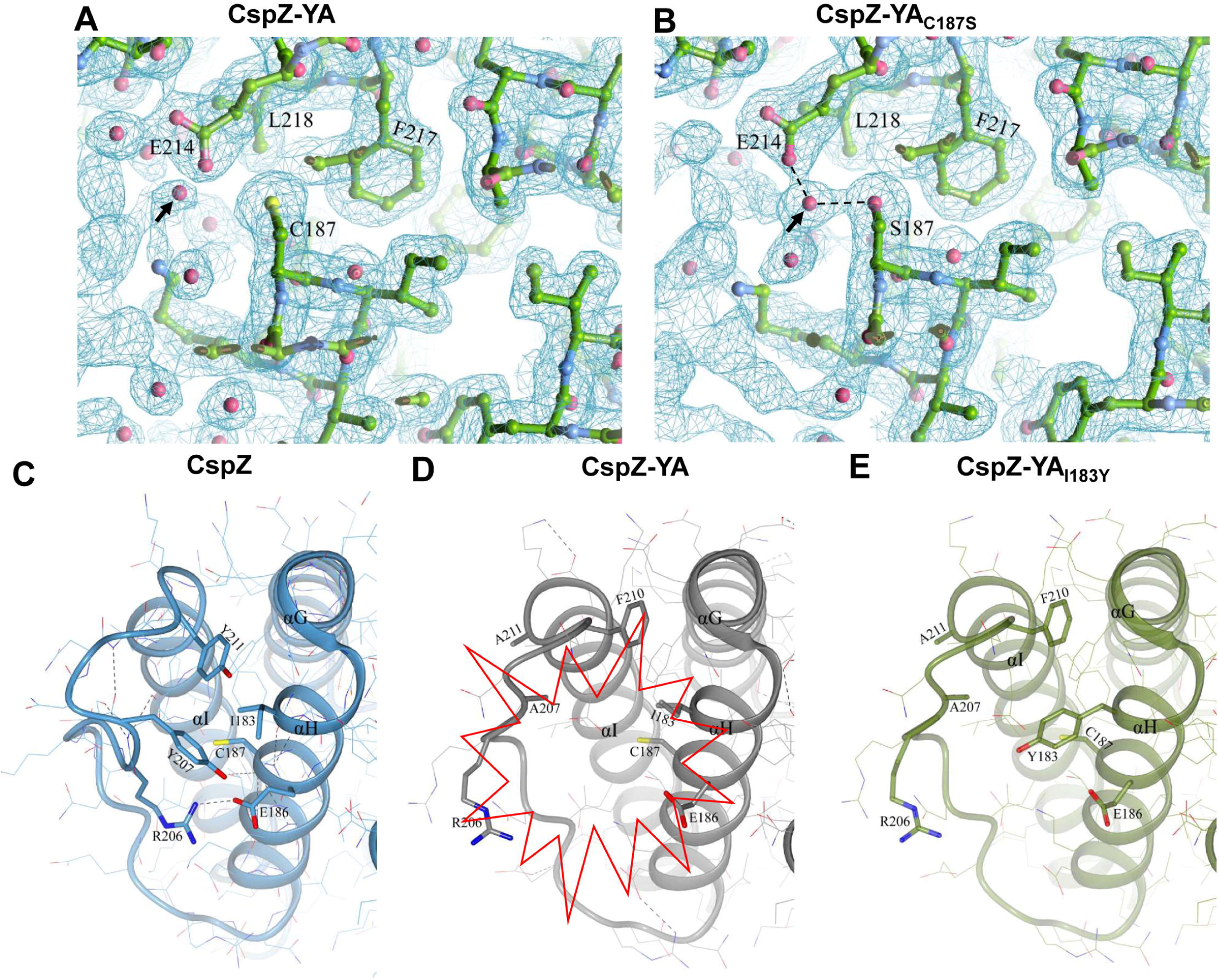
The comparison of CspZ, CspZ-YA, CspZ-YA_C187S_, and CspZ-YA_I183Y_ structures suggest the helix H-I interactions impacted by the C187S and I183Y mutagenesis. The structures here were obtained from CspZ (PDB ID 9F7I), CspZ-YA (PDB ID 9F1V), CspZ-YA_C187S_ (PDB ID 9F21), and AlphaFold predicted structure of CspZ-YA_I183Y_. (**A to B**) Shown is the 2Fo-Fc electron density map contoured at 1σ of the region around C187 in **(A)** CspZ-YA and S187 in **(B)** CspZ-YA_C187S._ The hydrogen bond formed between the water molecule with S187 and E214 were highlighted. **(C to E)** The crystal structures of **(C)** CspZ from *B. burgdorferi* B31, **(D)** CspZ-YA, and **(E)** the predicted structure of CspZ-YA_I183Y_ show the hydrophobic core accounting for helices G, H and I and the residues Y207, Y211, I183, and C187 in CspZ and the equivalent residues in CspZ-YA and CspZ-YA_I183Y_.

#### (iii) The I183Y and C187S mutations promoted the stability of the CspZ-YA epitopes recognized by CspZ-targeting, Lyme borrelia-killing monoclonal antibodies

The enhanced intramolecular interactions by I183Y and C187S mutagenesis raises a possibility of CspZ-YA_I183Y_ and CspZ-YA_C187S_ to have increasing stability. We thus examined whether CspZ-YA_I183Y_ and CspZ-YA_C187S_ have greater thermostability than CspZ-YA. We found indistinguishable Tm values between CspZ-YA_I183Y_ and CspZYA_C187S_ (61.87 and 62.72 °C, respectively) (**Fig. 6A and B, Table S3**). In contrast, the Tm-values of CspZ-YA were significantly lower, 57.58 and 58.46 °C for histidine tagged and untagged CspZ-YA, respectively, indicating a stability enhancement through mutagenesis (**Fig. 6A to B, Table S3**). We next examined the impact of I183Y and C187S mutagenesis on altering long-term stability of the protective epitopes in the CspZ-YA structures. We generated humanized, recombinant, and monoclonal CspZ-YA IgGs that contain the Fc region of human IgG1 and F(ab’)2 from 1139 or 1193, our two monoclonal CspZ-YA IgGs documented to eliminate Lyme borreliae [34]. The resulting humanized IgGs, namely 1139c and 1193c, were first confirmed for their ability to bind to CspZ-YA (**Fig. S5A and B**), block the FH-binding ability of CspZ (**Fig. S5C**), and promote lysis (**Fig. S5D**) and opsonophagocytosis of *B. burgdorferi* (**Fig. S5E**). We placed CspZ-YA_I183Y_, CspZYA_C187S_, or CspZ-YA (untagged and histidine tagged) at 4 or 37 °C for different period of time and then examined the ability of 1139c or 1193c to bind to each of these CspZ-YA proteins. We found all CspZ-YA proteins or variants previously incubated at 4 °C for 6- or 24-h or at 37 °C for 6-h displayed similar levels of recognition to these proteins prior to incubation (**Fig. 6C and D**). However, CspZ-YA but not CspZ-YA_I183Y_ and CspZ-YA_C187S_ previously incubated at 37 °C for 24-h had significantly lower levels of recognition, compared to those proteins prior to incubation (**Fig. 6C and D**), even though SE-HPLC did not indicate significant aggregation or degradation (results not shown). These findings demonstrated long-term stability enhancement of CspZ-YA proteins in the physiological temperature by I183Y and C187S mutagenesis, specifically on the structures that promote protective antibody induction.

**Figure 6.**
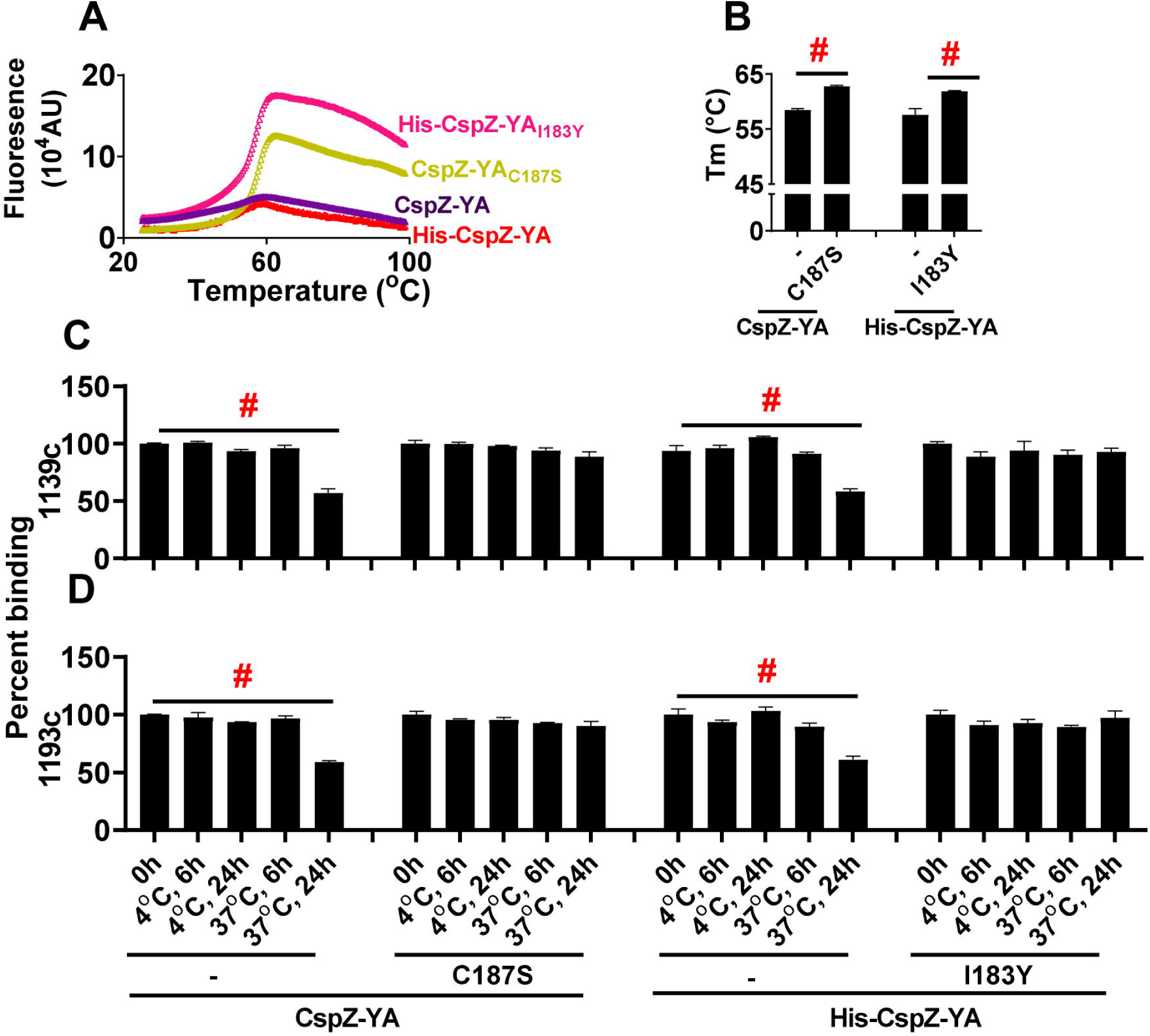
CspZ-YA_C187S_ and CspZ-YA_I183Y_ maintained the recognition by protective CspZ IgGs at higher temperature for longer period of time. **(A and B)** Untagged CspZ-YA or CspZ-YA_C187S_ (CspZ-YA_C187S_) or histidine tagged CspZ-YA (His-CspZ-YA) or CspZ-YA_I183Y_ (His-CspZ-YA_I183Y_) (10µM) in PBS buffer were subjected to the thermoshift assays described in the materials and methods. **(A)** Shown is the fluorescence intensities of each of the CspZ-YA proteins under the temperatures ranging from 25 to 99°C from one representative experiment. **(B)** The melting temperature (Tm) was extrapolated from the maximal positive derivative values of the fluoresces intensity (d(RFU)/dT) as bars. Data shown are the mean ± standard deviation of the Tm values for each of the CspZ-YA proteins from eight experiments. Statistical significance (p < 0.05, Kruskal Wallis test with the two-stage step-up method of Benjamini, Krieger, and Yekutieli) of differences in percent binding between groups are indicated (“#”). **(C and D)** One µg CspZ-YA, CspZ-YA_C187S_ (C187S), His-CspZ-YA, or His-CspZ-YA_I183Y_ (I183Y) was incubated at 4 or 37 °C for 6- or 24-h prior to being coated on microtiter plate wells. The microtiter plate wells immobilized with each of these proteins before incubation (0-h) were included as unincubated control. The ability of the CspZ monoclonal IgG, **(C)** 1139c or **(D)** 1193c, to recognize each of these CspZ-YA proteins were determined using ELISA in the section “Accelerated stability study” in Materials and Methods. The work was performed on four independent experiments (one replicate per experiment). Data are expressed as the percent binding, derived by normalizing the levels of bound 1139c or 1193c from the wells coated with each of the CspZ-YA proteins in different incubating conditions to that in the unincubated control. Data shown are the mean ± standard deviation of the percent binding of 1139c or 1193c from four experiments. Statistical significance (p < 0.05, Kruskal Wallis test with the two-stage step-up method of Benjamini, Krieger, and Yekutieli) of differences in percent binding between groups are indicated (“#”).

## DISCUSSION

Using native microbial surface antigens as vaccine targets presents multiple challenges, one of which is that native antigens are often not immunogenic, lacking the ability to provide protective immunity [4]. While other native antigens are immunogenic, conformational changes may occur after those antigens bind to their host ligands *in vivo* to promote pathogen invasion. Such conformational alteration of antigens could result in the potential of the induced antibodies to not constantly recognize pathogens and/or inhibit the pathogen invasion, decreasing vaccine efficacy [8, 42–45]. These difficulties were initially observed in some viral proteins but recently have also been reported in native antigens from non-viral pathogens, such as Lyme disease bacteria [32, 33]. One strategy to overcome these hurdles is through amino acid mutagenesis to force those antigens into certain structures that favor the induction of pathogen-killing antibodies ([34], for review paper, see [46]). For example, immunization with the native version of a Lyme borreliae FH-binding protein, CspZ, did not prevent Lyme disease infection [27, 32, 33, 47]. We previously generated CspZ-YA through the mutagenesis of Y207A and Y211A and found CspZ-YA to induce robust levels of borreliacidal antibodies that prevent the infection, suggesting a newly generated surface epitopes in CspZ-YA [32, 33]. In this study, our newly obtained high-resolution structure of CspZ-YA, paired with the previously resolved structure of CspZ-FH complex, provides evidence supporting the inability of CspZ-YA to bind to FH and exposed epitopes surrounding FH-binding sites [35]. Using CspZ as a model, our work here thus structurally demonstrated the potential mechanisms underlying the antigen engineering concept of unmasking the protective epitopes to promote vaccine efficacy.

The ability to maintain effective antibody titers determines the required immunization frequency of a vaccine. Engineering antigens to maintain preferred structures that can trigger long-lasting bactericidal antibodies remains an unresolved issue. Structure-based vaccine design is one of the recently developed strategies to promote the long-term antigen stability and vaccine efficacy [6, 7]. According to the existing high-resolution structures of an antigen, a series of amino acid residues are mutated with the goal to enhance antigen stability by promoting suitable intramolecular interactions [6, 7]. In this study, the mutagenesis of Y207A and Y211A reduced the hydrophobic interactions and created a cavity at the C-terminal H/I loop and helix H of CspZ-YA, destabilizing the protein’s conformation. Introduction of hydrophobic amino acid residues (i.e., isoleucine, leucine, valine, phenylalanine, and tyrosine) ideally would fill the cavity to maintain protein stability [48–50]. Such a “cavity filling” strategy has been examined by filling the hydrophobic cores of several viral antigens (e.g., stabilizing the perfusion structures of RSV F and the binding interface of HIV glycoproteins (gp120-gp41) [51, 52]). Here we included the I183Y mutation for cavity filling (**Fig. 5E**), elevating the efficacy of CspZ-YA-triggered bactericidal antibodies and preventing bacterial colonization and disease manifestations at lower immunization frequency. The stability enhancement by I183Y mutagenesis and the structure comparison between CspZ-YA_I183Y_ vs. CspZ-YA prove the concept of cavity filling at one of the first times in a bacterial vaccine antigen.

The other strategy of structure-based vaccine design is to manipulate disulfide bonds. Adding cysteine residues may lead to the formation of disulfide bonds, promoting intramolecular interaction and protein stability [53, 54]. However, the free cysteine residue in the antigens may also contribute to unwanted intermolecular interactions, resulting in protein aggregations [36–39]. Here, we attempted to test the impact of C187 on CspZ-YA in protein stability by replacing this residue with serine. We did not observe CspZ-YA and CspZ-YA_C187S_ forming apparent aggregates, but the reduced protective immunization frequency of CspZ-YA_C187S_ is correlated with the stability enhancement of this mutant protein. Our structural evidence further attributes such stability enhancement to the role of water molecule-coordinated helix H-I intramolecular interactions. In fact, sulfur had slightly greater van der Waals radius (∼1.8 Å), compared to oxygen (∼1.5 Å). Thus, the highly polar nature of serine together with a slightly smaller footprint of oxygen from S187 may be the cause of the water molecule positioning in the fashion to enhance hydrogen bond-mediated intramolecular interactions in CspZ-YA_C187S_ (**Fig. 5A and B**). Overall, this work provides the structural evidence underlying the cysteine mutagenesis-mediated stability enhancement. Further, as mutations of I183 and C187 are both located on helix H and facilitating helix H-I interactions, these findings may raise an intriguing possibility to engineer other amino acids on helix H or I in promoting intramolecular interactions (specifically, helix H-I interactions), which is worth for further investigations.

We found our CspZ-targeted and protective monoclonal antibodies to recognize CspZ-YA_I183Y_ and CspZ-YA_C187S_, better than CspZ-YA at a higher temperature for a longer period of time (i.e., 37°C for 24-h). As mammalian body temperatures stay consistent at 37°C, both mutations would allow CspZ-YA to persist in the designated structures to promote the continuous production of resulting protective antibodies, thus suitable as vaccines for human or other mammal uses. Additionally, I183 and C187 are both located on helix H. The structural comparison of CspZ-YA_C187S_ and CspZ-YA_I183Y_ with their parental CspZ-YA correlated I183Y- and C187S-mediated stability enhancement with the facilitation of favorable helix H-I interactions on CspZ-YA (**Fig. 5**). Although I183 and C187 are not located on or surrounding the FH-binding interface of CspZ, the N-terminus of helix H and the loop H/I are within and immediately adjacent to the FH-binding sites [35](Brangulis et al. unpublished). Therefore, it is conceivable that strengthening helix H-I interactions of CspZ-YA vaccines stabilize the structures of the protective epitopes within or adjacent to the FH-binding interface of CspZ, triggering greater levels of protective antibodies. Testing this possibility would require the elucidation of the high-resolution complexed structure of CspZ-YA and those CspZ-targeted and protective monoclonal antibodies (i.e., 1139c and 1193c), requiring further investigations.

The more stable structures suggested by structure-based vaccine design would also potentially aid the vaccine production and ease the required conditions for transportation and storage, promoting the commercialization plan [55]. In this study, we mutagenized CspZ-YA as a model to test the concept of structure-based vaccine design in decreasing the minimal immunization frequency that allows protectivity. The results elucidate the mechanisms underlying such a concept using a Lyme disease subunit vaccine as a model. Additionally, the fact that CspZ-YA_I183Y_ or CspZ-YA_C187S_ provides lower protective immunization frequency than OspA offers the opportunity to use these mutated proteins as vaccines to overcome the need for constant immunization for OspA-targeted vaccines. Finally, the breakthrough of vaccine design sparked off by recent pandemics underscores the importance of structure-guided approaches for efficacy optimization of vaccines. This concept-proof study thus provides mechanistic insights into structural-based vaccine design and illustrates the possibility of revisiting the previously tested but inefficacious antigens. This work would hopefully facilitate the establishment of a pipeline for vaccine design that can be extended to combat other newly emerging infectious diseases.

## MATERIALS AND METHODS

### Ethics Statement

All mouse experiments were performed in strict accordance with all provisions of the Animal Welfare Act, the Guide for the Care and Use of Laboratory Animals, and the PHS Policy on Humane Care and Use of Laboratory Animals. The protocol (Docket Number 22-451) was approved by the Institutional Animal Care and Use Agency of Wadsworth Center, New York State Department of Health. All efforts were made to minimize animal suffering. This study also involves secondary use of deidentified archival patient sera collected in previous studies and was approved by the Institutional Review Board (IRB) of New York State Department of Health and Baylor College of Medicine under protocol 565944-1 and H-46178, respectively. Analysis of deidentified patient data was carried out under a waiver of consent.

### Mouse, ticks, bacterial strains, hybridoma, and human serum samples

Four-week-old, female C3H/HeN mice were purchased from Charles River (Wilmington, MA, USA). Although such an age of the mice has not reached sexual maturity, the under development of immune system in this age of mice would allow such mice to be more susceptible to Lyme borreliae infection, increasing the signal to noise ratio of the readout. That will also provide more stringent criteria to define the protectivity. BALB/c C3-deficient mice were from in-house breeding colonies [56] and *Ixodes scapularis* tick larvae were obtained from BEI Resources (Manassas, VA). *Escherichia coli* strain BL21(DE3) and derivatives were grown at 37°C or other appropriate temperatures in Luria-Bertani broth or agar, supplemented with kanamycin (50µg/mL). *B. burgdorferi* strain B31-A3 were grown at 33°C in BSK II complete medium [57]. Cultures of *B. burgdorferi* B31-A3 were tested with PCR to ensure a full plasmid profile before use [58, 59]. Hybridoma that produce the monoclonal antibodies #1139c or #1193c were cultivated in RPMI 1640 medium containing 10% FBS at 37°C with 5% of CO_2_. Thirty-eight deidentified two-tiered positive human serum samples were obtained from New York State Department of Health. These serum samples were previously collected from humans that were tested positive in two-tiered assays, which is the serological definition of Lyme disease infection [60]. The negative control human sera **were** collected from 10 individuals residing in a non-endemic area **for** Lyme disease.

### Cloning, expression and purification of OspA, CspZ, CspZ-YA, and CspZ-YA-derived mutant proteins

The DNA encoding histidine tagged CspZ, CspZ-YA and CspZ-YA-derived mutant proteins (**Table S4**) was codon-optimized based on *E. coli* codon usage preference and synthesized by Synbiotech (Monmouth Junction, NJ), followed by subcloning into the pET28a using BamHI/SalI restriction sites. These plasmids were transformed into *E. coli* BL21 (DE3). The DNA encoding untagged CspZ and its derived mutant proteins were codon-optimized based on *E. coli* codon usage preference, synthesized and subcloned into the pET41a using NdeI/XhoI sites by GenScript (Piscataway, NJ). These plasmids were transformed into *E. coli* BL21 (DE3). The recombinant protein expression was induced with 1 mM Isopropyl-β-D-1-thiogalactopyranoside (IPTG). Once expression was confirmed, the clone with the highest expression for each construct was selected to create glycerol seed stocks. The generation of histidine-tagged CspZ and this protein-derived mutant proteins is described previously [61]. To purify the untagged CspZ-YA, and CspZ-YA_C187S_, we followed the procedure as described[61]. Because lipidation is required for recombinant OspA proteins to protect mice from Lyme disease infection [62, 63], the lipidated OspA was included as a control. To generate the lipidated OspA, the previous process was followed [63]. For structural studies, the encoding regions of CspZ-YA and CspZ-YA_C187S_ were cloned into the pETm-11 expression vector containing an N-terminal 6xHis tag, followed by a tobacco etch virus (TEV) protease cleavage site. Both proteins were expressed in *E. coli* BL21 (DE3) and purified by affinity chromatography as described previously for CspZ [64].

### Generation of humanized CspZ-YA antibodies, #1139c and #1193c

Protective mouse monoclonal antibodies (mAbs) 1139 and 1193 against CspZ were developed previously [34]. These two mAbs were further humanized using the service provided by GenScript Probio (Piscataway, NJ). Briefly, DNA sequencing was performed using the hybridoma to identify the gene coding the variable domain of mAbs 1139 and 1193. Such genes were then grafted with the one coding for human IgG1. The two humanized chimeric mAbs (1139c and 1193c) were then transiently produced in CHO cells, followed by purification with Protein A affinity chromatography.

### Circular dichroism (CD) spectroscopy

CD analysis was performed on a Jasco 810 spectropolarimeter (Jasco Analytical Instrument, Easton, MD) under nitrogen. CD spectra were measured at room temperature (RT, 25°C) in a 1mm path length quartz cell. Spectra of each of the CspZ-YA proteins (10μM) were recorded in phosphate based saline buffer (PBS) at RT, and three far-UV CD spectra were recorded from 190-250 nm in 1 nm increments for far-UV CD. The background spectrum of PBS without proteins was subtracted from the protein spectra. CD spectra were initially analyzed by the software Spectra Manager Program (Jasco). Analysis of spectra to extrapolate secondary structures was performed using the K2D3 analysis programs [65].

### Mouse immunization and infection

C3H/HeN Mice were immunized as described, with slight modifications [33]. Fifty µl of PBS (control) or 25 µg of untagged or histidine tagged CspZ-YA or its mutant proteins, or untagged, lipidated OspA in 50µl of PBS was thoroughly mixed with 50µl TiterMax Gold adjuvant (Norcross, GA, USA), resulting in total 100µl of the inoculum. This inoculum was introduced into C3H/HeN mice subcutaneously once at 0 days post initial immunization (dpii), twice at 0 and 14 dpii, or three times at 0, 14, and 28 dpii (**Fig. 1**). At 14 days post last immunization (dpli), blood was collected via submandibular bleeding to isolate serum for the determination of ability in recognizing CspZ, CspZ-YA and the mutant proteins derived from CspZ-YA, as described in the sections of “ELISAs” and “Borreliacidal assays”, respectively (**Fig. 1**). At 7 dpli, *B. burgdorferi* B31-A3-infected flat nymphs were placed in a chamber on the immunized or PBS-inoculated C3H/HeN mice as described (**Fig. 1**)[47]. Five nymphs were allowed to feed to repletion on each mouse, and a subset of nymphs was collected pre- and post-feeding. At 21 dpli, tick bite sites of skin, bladder, knees, and heart were collected to determine the bacterial burdens, and ankles were also collected at 21 dpli to determine the severity of arthritis described in the section “Quantification of spirochete burdens and histological analysis of arthritis. (**Fig. 1**).” At this time point, blood was also collected via cardiac puncture bleeding to isolate serum for the determination of seropositivity described in the section “ELISAs” (**Fig. 1**).

For the mice inoculated with humanized monoclonal IgGs, C3H/HeN mice were immunized as described, with slight modifications [34]. Basically, C3H mice were intraperitoneally inoculated with irrelevant human IgG (control), #1139c or #1193c (1 mg/kg) (**Fig. S6A**). Five mice per group were used in this study. At 24 hours after inoculation, five nymphs carrying *B. burgdorferi* strain B31-A3 were allowed to feed to repletion on each mouse, and a subset of nymphs was collected pre- and post-feeding as described [33, 56]. Mice were sacrificed at 21 days post feeding (dpf) to collect the biting site of skin, bladder, knees, and heart to determine the bacterial burdens described in the section “Quantification of spirochete burdens and histological analysis of arthritis (**Fig. S6A**).” Blood was also collected via cardiac puncture bleeding to isolate sera for the determination of seropositivity described in the section “ELISAs” (**Fig. S6A**).

### ELISAs

To measure the titers of anti-CspZ IgG in the serum samples (**Fig. S2**), one µg of histidine-tagged CspZ was coated on ELISA plate wells as described [33]. To determine the ability of anti-CspZ IgG in the sera to recognize CspZ-YA, and CspZ-YA_I183Y_, and CspZ-YA_C187S_ (**Fig. 4A**), each of these proteins with histidine tags (1 µg) was coated on ELISA plate wells as in the same fashion. The procedures following the protein coating are as described previously [33]. For each serum sample, the maximum slope of optical density/minute of all the dilutions of the serum samples was multiplied by the respective dilution factor, and the greatest value was used as arbitrary unit (A.U) to represent the antibody titers for the experiment to obtain anti-CspZ IgG (**Fig. S2**) or the ability of the anti-CspZ IgG in the sera to recognize CspZ-YA and different CspZ-YA mutant proteins (**Fig. 4A**). The quality of the correlation for the ability of those CspZ IgG in recognizing CspZ-YA vs. CspZ-YAC187S, CspZ-YA vs. CspZ-YAI183Y, or CspZ-YAC187S vs. CspZ-YAI183Y was determined by the R and P values of Spearman analysis, which was calculated using dose-response stimulation fitting in GraphPad Prism 9.3.1.

Additionally, the seropositivity of the mice after infection with *B. burgdorferi* was determined by detecting the presence or absence of the IgGs that recognize C6 peptides (**Fig. 3A, S3A, and S6A**). This methodology has been commonly used for human Lyme disease diagnosis [66] and performed as described in our previous work [34]. For each serum sample, the maximum slope of optical density/minute of all the dilutions was multiplied by the respective dilution factor, and the greatest value was used as representative of anti-C6 IgG titers (arbitrary unit (A.U.)). The seropositive mice were defined as the mice with the serum samples yielding a value greater than the threshold, the mean plus 1.5-fold standard deviation of the IgG values derived from the uninfected mice.

We also determined the ability of #1139c or #1193c to prevent FH from binding to CspZ (**Fig. S5C**), which was performed as described previously with modifications [34]. Basically, each ELISA microtiter well was coated with one µg of histidine-tagged CspZ. After being blocked with 5% BSA in PBS buffer, the wells were incubated with PBS (control) or serially-diluted irrelevant human IgG (Human IgG isotype control, Sigma-Aldrich, St. Louis, MO) #1139c or #1193c (0.4 nM, 0.8 nM, 1.6 nM, 3.125 nM, 6.25 nM, 12.5 nM, 25 nM, 50 nM) followed by being mixed with 500 nM of human FH. Sheep anti-human FH (1:200×, ThermoFisher; Waltham, MA) and then donkey anti-sheep HRP (1:2000×, ThermoFisher) were added, and the levels of FH binding were detected by ELISA as described previously [34]. Data were expressed as the proportion of FH binding from serum-treated to PBS-treated wells. The 50% inhibitory concentration (IC_50_) (the inlet figure of **Fig. S5C**), representing the IgG concentration that blocks 50% of FH binding, was calculated using dose-response stimulation fitting in GraphPad Prism 9.3.1.

### Borreliacidal assays

The ability of serum samples (**Fig. 2B to G**) or humanized monoclonal CspZ IgG (#1139c and #1193c, **Fig. S5D**) to eradicate *B. burgdorferi* B31-A3 was determined as described with modifications [32, 33]. Briefly, the sera collected from mice immunized with different CspZ-YA proteins at different immunization frequency were heat-treated to inactivate complement. Each of these serum samples or #1139c or #1193c was serially diluted, and mixed with complement-preserved guinea pig serum (Sigma-Aldrich) or heat-inactivated guinea pig serum (negative control). After adding the strain *B. burgdorferi* B31-A3, the mixture was incubated at 33°C for 24 hours. Surviving spirochetes were quantified by directly counting the motile spirochetes using dark-field microscopy and expressed as the proportion of serum-treated to untreated Lyme borreliae. The 50% borreliacidal activities (BA_50_), representing the serum dilution rate (for the serum samples in **Fig. 2B to G**) or the concentration of IgGs (for #1139c and #1193c in **Fig. S5D**) that kills 50% of spirochetes, was calculated using dose-response stimulation fitting in GraphPad Prism 9.3.1.

### Quantification of spirochete burdens and histological analysis of arthritis

DNA was extracted from the indicated mouse tissues to determine the bacterial burdens (**Fig. 3B to F, S3B to F and H to L, and S6C to G**), using quantitative PCR analysis as described [33]. Note that spirochete burdens were quantified based on the amplification of *recA* using the forward and reverse primers with the sequences as GTGGATCTATTGTATTAGATGAGGCTCTCG and GCCAAAGTTCTGCAACATTAACACCTAAAG, respectively. The number of *recA* copies was calculated by establishing a threshold cycle (Cq) standard curve of a known number of *recA* gene extracted from strain B31-A3, and burdens were normalized to 100 ng of total DNA. For the ankles that were applied to histological analysis of Lyme disease-associated arthritis (**Fig. 3G**), the analysis was performed as described [33]. The image was scored based on the severity of the inflammation as 0 (no inflammation), 1 (mild inflammation with less than two small foci of infiltration), 2 (moderate inflammation with two or more foci of infiltration), or 3 (severe inflammation with focal and diffuse infiltration covering a large area).

### Crystallization and structure determination

Initial crystallization trials of CspZ-YA and CspZ-YA_C187S_ were performed in 96-well sitting drop crystallization plates (SWISSCI AG, Neuheim, Switzerland), using sparse-matrix screens JCSG+ and Structure Screen 1&2 from Molecular Dimensions (Newmarket, UK). Tecan Freedom EVO100 workstation (Tecan Group, Männedorf, Switzerland) was used to set up the plates by mixing 0.4 μl of protein with 0.4 μl of precipitant. After initial crystal hits, the corresponding crystallization conditions were optimized by varying the quantities of the components in the precipitant solution to obtain crystals suitable for harvesting. Diffraction data for CspZ-YA was collected from crystals grown in 0.2 M ammonium acetate, 0.1 M sodium citrate (pH 6.5) and 30% PEG 4000, but for CspZ-YA_C187S_ grown in 2.2 M ammonium citrate, 0.1 M HEPES (pH 7.5) and 2% PEG 400. Before harvesting and storing the crystals in liquid nitrogen, crystals for CspZ-YA were subjected to cryoprotectant made of the precipitant solution with additional 10% glycerol. Diffraction data for CspZ-YA were collected at the Diamond Light Source (Oxfordshire, UK) beamline I03 but the data for CspZ-YA_C187S_ at the MX beamline instrument BL 14.1 at Helmholtz-Zentrum (Berlin, Germany) [67]. Reflections were indexed by XDS and scaled by AIMLESS from the CCP4 suite [68, 69]. Initial phases for CspZ-YA and CspZ-YA_C187S_ were obtained by molecular replacement using Phaser [70], with the crystal structure of *B. burgdorferi* CspZ as a search model (PDB ID 4CBE). After molecular replacement, the protein models were built automatically in BUCCANEER [71]. The crystal structures were improved by manual rebuilding in COOT [72]. Crystallographic refinement was performed using REFMAC5 [73]. A summary of the data collection, refinement and validation statistics for CspZ-YA and CspZ-YA_C187S_ are given in **Table S2.**

### Protein 3D structure prediction using AlphaFold

AlphaFold v2.0 [74] was used to predict the 3D structure for CspZ-YA_I183Y_ as described previously for *B. burgdorferi* PFam12 family proteins [75].

### Surface Plasmon Resonance (SPR)

Interactions of CspZ-YA with #1139c or #1193c were analyzed by SPR using a Biacore T200 (Cytiva, Marlborough, MA). Ten micrograms of #1139c or #1193c were conjugated to a Sensor Chip Protein G (Cytiva) by flowing each of these IgGs at the flow rate at 10μl/min, 25 °C through that chip using PBS as the buffer. For quantitative SPR experiments, 10µL of increasing concentrations (0, 15, 31.25, 62.5, 125, 250, 500 nM) of CspZ-YA were injected into the control cell and the flow cell immobilized with #1139c or #1193c at 10μl/min, 25°C. To obtain the kinetic parameters of the interaction, sensogram data were fitted by means of BIAevaluation software version 3.0 (GE Healthcare), using the one step biomolecular association reaction model (1:1 Langmuir model), resulting in optimum mathematical fit with the lowest Chi-square values.

### Phagocytosis assays

The phagocytosis assays were performed as described previously with modifications [76]. *B. burgdorferi* B31-A3 were labeled with carboxyfluorescein diacetate succinimidyl ester (CFSE, Invitrogen) as described in vendor’s manual. Basically, the suspension of spirochetes (10^7^) in BSK II media without rabbit sera, gelatin, and BSA was incubated with 3.3 μM of CFSE at room temperature for 10 minutes. To prepare the antibody-treated sera, normal or heat-inactivated human sera that are determined negative to anti-C6 IgGs were incubated with CFSE labeled spirochetes (10^7^ bacteria) in the presence of #1139c, #1193c, or irrelevant human IgG (Human IgG isotype control, Sigma-Aldrich) at room temperature for 10 minutes. Such spirochete suspension was then mixed with freshly isolated human neutrophils (PMNs) from a blood donor iQBioscience (Alameda, CA) at the ratio of 25 to 1 and shaking at 37 °C, 50 rpm for 10 minutes. For each sample, the bacteria-PMNs mixture incubated on ice for 10 minutes immediately after mixing was included as control. Phagocytosis was stopped by transferring the bacteria-PMN mixtures to ice-cold Fluorescence-Activated Cell Sorting (FACS) buffer (PBS supplemented with 0.5% bovine serum albumin (BSA), 0.01% NaN3 and 0.35 mM EDTA) and stored at 4 °C. Samples continually kept on 4 °C were used as a control. PMNs were then washed suspended with ice-cold FACS-buffer prior to be applied to a FACSCalibur flow cytometer (Beckton Dickinson). The phagocytosis index of each sample was calculated as mean fluorescence intensity (MFI)×percentage (%) positive cells) at 37°C minus (MFI×% positive cells) at 4 °C. Each sample were performed in seven replicates in two different events.

### Fluorescence-based thermal shift assays

Ten µM of indicated wild-type and mutant CspZ-YA proteins was applied to 7500 Fast Real-Time PCR System (Thermo Scientific) with a temperature range of 25–95 °C. All reactions were in 20 µl of the final volume in 96-well plates using Protein Thermal Shift™ Dye Kit (ThermoFisher Scientific) at 1:1,000 dilution in PBS buffer. The protein-unfolding concentration (Tm) were extrapolated by obtaining the temperature with maximal positive derivative values of the fluorescence intensity using the 7500 Fast Real-Time PCR System software (Thermo Scientific).

### Accelerated stability study

One µg of untagged CspZ-YA or CspZYA_C183S_, or histidine-tagged CspZ-YA or CspZYA_I183Y_ was incubated at 4 or 37°C for 6- or 24-h prior to be coated on ELISA plate wells as described [33]. The ELISA plate wells immobilized with untagged or histidine-tagged CspZ-YA before incubation were included as control. After blocking those plate wells by PBS with tween 20 as described [33], #1139c or #1193c (1µM), was added to those wells, and the levels of binding between each of these antibodies with CspZ-YA proteins were determined by ELISA as described in the section “ELISAs.” Data were expressed as the proportion of #1139c- or #1193c-binding from the ELISA plate wells immobilized with the CspZ-YA proteins incubated at different conditions to those with the control wells.

### Statistical analyses

Significant differences were determined with a Kruskal-Wallis test with the two-stage step-up method of Benjamini, Krieger, and Yekutieli [77], two-tailed Fisher test (for seropositivity in Fig. 3A, S3A, and S6B)[78], or Spearman analysis (for correlation analysis in Fig. 4B to D and F to H) [79], using GraphPad Prism 9.3.1. A p-value < 0.05 was used to determine significance.

## ACKNOWLEDGEMENTS

The authors thank Patricia Rosa and John Leong for providing *B. burgdorferi* strain B31-A3. They also thank Klemen Strle for valuable advice. The authors also thank the Wadsworth Animal Core for assistance with Animal Care and Leslie Eisele and Renjie Song of Wadsworth Biochemistry and Immunology Core for CD spectroscopy, SPR, and flow cytometry, and Susan Wong from Wadsworth Diagnostic Immunology Laboratory to provide human sera. Diffraction data for *B. burgdorferi* CspZ-YA were collected on Diamond Light Source (Oxfordshire, UK) on I03 beamline for beamtime MX35587-1, but for CspZ-YA_C187S_ on beamline BL14.1 at the BESSY II electron storage ring operated by the Helmholtz-Zentrum (Berlin, Germany). This work was supported by NIH grant R01AI181746 (for A.L.M., Y.L.), R44AI152954 (for J.M., A.L.M., A.W., Y.L.), R21AI144891 and the U.S. Department of Defense, Congressionally Directed Medical Research Programs, Grant Number W81XWH-20-1-0913 (Y.C., A.L.M., T.A.N., M.E.B, W.H.C., Y.L. R.T.K, ZL), and R01AI154542 (for X.Y., U.P.). The funders had no role in study design, data collection, interpretation, or the decision to submit the work for publication. Y. L. is the inventor on U.S. patent application no. US11771750B2 (“Composition and method for generating immunity to *Borrelia burgdorferi*”). The remaining authors declare no competing interests.

