## Supplemental Figure Legends, and Tables for "Mechanistic insights into structure-based design of a Lyme disease subunit vaccine"

### SUPPLEMENTAL MATERIALS

### SUPPLEMENTAL FIGURE LEGENDS

**Figure S1. CD spectra demonstrate no impacts of secondary structures by mutating indicating amino acids of CspZ-YA.** Far-UV CD analysis of **(A)** untagged CspZ-YA and CspZ-$YA_{C187S}$ (CspZ-$YA_{C187S}$), and **(B)** histidine tagged CspZ-YA (His-CspZ-YA) and the mutant proteins derived from this protein. The molar ellipticity, Φ, was measured from 190-250nm for 10μM of each protein in PBS.

**Figure S2. Immunization of CspZ-YA and its mutant proteins triggered indistinguishable levels of antibodies against CspZ.** Sera were collected at 14dpli from C3H/HeN mice immunized **(A)** once, **(B)** twice, or **(C)** three times in the fashion as described in Fig. 1. These mice were immunized with PBS (control) or untagged CspZ-YA or its derived mutant protein, or histidine tagged CspZ-YA (His-CspZ-YA), or its derived mutant proteins (Six mice for CspZ-YA- or CspZ-$YA_{C187S}$-immunized mice whereas five mice for the rest of immunization groups of mice). The levels of total IgG against CspZ were determined using quantitative ELISA. Data shown are the geometric mean ± geometric standard deviation of the titers of anti-CspZ antibodies from five mice per group. Asterisks indicate the statistical significances ($p < 0.05$, Kruskal Wallis test with the two-stage step-up method of Benjamini, Krieger, and Yekutieli) of differences in antibody titers relative to the sera from PBS-inoculated mice.

**Figure S3. Immunization once or three times with CspZ-YA, CspZ-$YA_{C187S}$, or CspZ-$YA_{I183Y}$ showed indistinguishable protectivity for seroconversion and borrelial tissue colonization.** Five PBS- or lipidated OspA (OspA)-, or histidine tagged CspZ-YA (His-CspZ-YA)- or CspZ-

$YA_{I183Y}$ (I183Y)-, or six untagged CspZ-YA- or CspZ-YAC187S (C187S)-immunized C3H/HeN mice that were immunized **(A to F)** once or **(G to L)** three times by indicated proteins in the fashion as described in Fig. 1. At 21 days post last immunization, these mice were then fed on by nymphs carrying *B. burgdorferi* B31-A3. Mice inoculated with PBS that are not fed on by nymphs were included as an uninfected control group (uninfect.). **(A and G)** Seropositivity was determined by measuring the levels of IgG against C6 peptides in the sera of those mice at 42 days post last immunization using ELISA. The mouse was considered as seropositive if that mouse had IgG levels against C6 peptides greater than the threshold, the mean plus 1.5-fold standard deviation of the IgG levels against C6 peptides from the PBS-inoculated, uninfected mice (dotted line). The number of mice in each group with the anti-C6 IgG levels greater than the threshold (seropositive) is shown. Data shown are the geometric mean ± geometric standard deviation of the titers of anti-C6 IgG. Statistical significances ($p < 0.05$, Kruskal-Wallis test with the two-stage step-up method of Benjamini, Krieger, and Yekutieli) of differences in IgG titers relative to (*) uninfected mice are presented. **(B to F, H to L)** *B. burgdorferi* (*Bb*) burdens at **(B and H)** nymphs after when feeding to repletion or **(C and I**) the tick feeding site ("Bite Site"), **(D and J)** bladder, **(E and K)** heart, and **(F and L)** knees, were quantitatively measured at 42 days post last immunization, shown as the number of *Bb* per 100ng total DNA.  Data shown are the geometric mean ± geometric standard deviation of the spirochete burdens from each group of mice. Asterisks indicate the statistical significance ($p < 0.05$, Kruskal Wallis test with the two-stage step-up method of Benjamini, Krieger, and Yekutieli) of differences in bacterial burdens relative to uninfected mice.

**Figure S4. More than 90% of Lyme disease human patients develop CspZ antibodies.** Sera from 38 patients with seropositive for Lyme disease infection (Two tier pos.; Positive in Two tier test) were determined for the titers of antibodies that recognize untagged CspZ using ELISA, as described in the section "ELISA" of the Materials and Methods. Ten serum samples from humans residing in non-endemic area of Lyme disease were included as negative control (Neg. ctrl.) and to set up the threshold value of titers that can be used to determine CspZ antibody positivity. That threshold value was mean 1.5-folds of standard deviation extrapolated from the values of negative control human sera. Thirty six out of 38 serum samples (94.7%) yield greater anti-CspZ IgG titers than the threshold values and was thus considered positive for CspZ antibodies. Shown is the geometric mean ± geometric standard deviation of the titers. Statistical significance ($p < 0.05$, Kruskal Wallis test with the two-stage step-up method of Benjamini, Krieger, and Yekutieli) of differences in CspZ-IgG titers between groups are indicated ("#").

**Figure S5. The humanized monoclonal antibodies 1139c and 1193c efficiently recognize CspZ-YA, prevent human FH-binding, and promote lysis and opsonophagocytosis of *B. burgdorferi*. (A and B)** The humanized monoclonal antibody **(A)** #1139c or **(B)** #1193c was flowed over the chip surface, conjugated with indicated untagged CspZ-YA. Binding was measured in response units (R.U.) by surface plasmon resonance. Shown is the mean ± standard deviation of the $k_{on}$, $k_{off}$, and $K_D$ values extrapolated from three experiments. One represented experiment is shown in this panel. **(C)** The monoclonal antibody #1139c or #1193c, or irrelevant human IgG (control, irr. hIgG) at indicated concentrations or PBS (control, data not shown) was added into the CspZ-coated ELISA plate wells. Each of those wells was then incubated with human FH, and the levels of bound FH were quantified using sheep anti-human FH and goat anti-sheep HRP IgG as primary and secondary antibodies, respectively. The work was performed on three independent experiments; within each experiment, samples were run in triplicate. Data are expressed as the percent human FH binding, derived by normalizing the levels of bound human FH from IgG-treated wells to that from PBS-treated wells. Data shown are the mean ± SEM of the percent human FH binding from three replicates. Shown is one representative experiment. The concentrations of the IgG to inhibit 50% of human FH bound by CspZ ($IC_{50}$) was obtained from curve-fitting and shown in the inlet figure. The $IC_{50}$ values are shown as the mean ± SD of from three experiments. **(D)** The monoclonal antibody #1139c or #1193c, or irrelevant human IgG (control, irr. hIgG) or PBS (control, data not shown) were serially diluted as indicated, and mixed with guinea pig complement and *B. burgdorferi* strains B31-A3 ($5 \times 10^5$ cells $ml^{-1}$). After incubated for 24 hours, surviving spirochetes were quantified from three fields of view for each sample using dark-field microscopy. The work was performed on three independent experiments. The survival percentage was derived from the proportion of IgG-treated to PBS-treated spirochetes. Shown is one representative experiment, and in that experiment, the data points are the mean ± SEM of the survival percentage from three replicates. The 50% borreliacidal activity of each IgGs ($BA_{50}$), representing the IgG concentrations that effectively killed 50% of spirochetes, was obtained and extrapolated from curve-fitting and shown in the inlet figure. The $BA_{50}$ values are shown as the mean ± SD of from three experiments.

**Figure S6. The humanized monoclonal antibodies 1139c and 1193c prevent seroconversion and tissue colonization caused by *B. burgdorferi* B31-A3 infection. (A)** Timeframe of the IgG inoculation and *B. burgdorferi* infection. **(B to G)** Five C3H/HeN mice were inoculated with the monoclonal antibody #1139c or #1193c, or irrelevant human IgG (control, irr. hIgG) at the dose of 1 mg/kg. At 24 hours after IgG inoculation, these mice were fed on by *I. scapularis* nymphs carrying *B. burgdorferi* B31-A3 (*Bb* B31-A3). An additional five mice inoculated with PBS but not fed on by ticks were included as the control (Uninfect.). The tissues were collected from those mice at 4 days post nymph feeding. Spirochete burdens at **(B)** the tick feeding site ("Bite Site"), **(C)** bladder, **(D)** heart, and **(E)** knees were quantitatively measured at 21 dpf, shown as the number of spirochetes per 100ng total DNA. Data shown are the geometric mean ± geometric standard deviation of the spirochete burdens from five mice per group. Statistical significances ($p < 0.05$, Kruskal-Wallis test with the two-stage step-up method of Benjamini, Krieger, and Yekutieli) of differences in bacterial burdens relative to (*) uninfected mice are presented.

### SUPPLEMENTAL TABLES

**Table S1. Statistics for Data and Structure Quality.**

| **Dataset** | **CspZ-YA** | **CspZ-YA$_{C187S}$** |
|---|---|---|
| **Space group** | $P2_12_12_1$ | $P2_12_12_1$ |
| **Unit cell dimensions** | | |
| **a (Å)** | 31.47 | 31.55 |
| **b (Å)** | 41.55 | 41.68 |
| **c (Å)** | 162.56 | 162.81 |
| **Wavelength (Å)** | 0.9762 | 0.9184 |
| **Resolution (Å)** | 162.56-1.90 | 41.68-2.00 |
| **Highest resolution bin (Å)** | 1.94-1.90 | 2.05-2.00 |
| **No. of reflections** | 231969 | 189839 |
| **No. of unique reflections** | 17550 | 15299 |
| **Completeness (%)** | 99.5 (100.0) | 99.9 (100.0) |
| **$R_{merge}$** | 0.09 (0.46) | 0.09 (0.38) |
| **$I/\sigma$ ($I$)** | 15.5 (4.8) | 19.3 (6.2) |
| **Multiplicity** | 13.2 (13.5) | 12.4 (13.5) |
| **Refinement** | | |
| **$R_{work}$** | 0.190 (0.245) | 0.248 (0.204) |
| **$R_{free}$** | 0.241 (0.308) | 0.335 (0.337) |
| **Average B-factor (Å$^2$)** | | |
| **Overall** | 28.4 | 29.8 |
| **From Wilson plot** | 17.0 | 17.5 |
| **No. of atoms** | | |
| **Protein** | 1743 | 1743 |
| **Water** | 199 | 170 |
| **RMS deviations from ideal** | | |
| **Bond lengths (Å)** | 0.009 | 0.008 |
| **Bond angles (°)** | 1.530 | 1.457 |
| **Ramachandran outliers (%)** | | |
| **Residues in most favored regions (%)** | 94.42 | 94.88 |
| **Residues in allowed regions (%)** | 4.65 | 4.19 |
| **Outliers (%)** | 0.93 | 0.93 |

Values in parentheses are for the highest resolution bin.

**Table S2. $BA_{50}$ values of the sera derived from mice immunized with CspZ-YA proteins in different immunization frequency**

| $BA_{50}$[a] | CspZ-YA[b] | | His-CspZ-YA[c] | | | | | | | | |
|---|---|---|---|---|---|---|---|---|---|---|---|
| Immunization frequency | - | C187S | - | T67P | I80T | F105P | I115T | K136E | V142M | I183Y | G193M |
| 1[d] | 25±1.6 | 19±1.2 | 26±1.1 | 20±1.5 | 24±1.6 | 29±1.0 | 25±1.2 | 17±1.2 | 21±1.3 | 29±1.3 | 27±1.1 |
| 2[d] | 43±1.3 | 204±1 | 43±1.5 | 45±1.1 | 56±1.1 | 53±1.0 | 53±1.3 | 41±1.1 | 35±1.3 | 164±1.0 | 48±1.0 |
| 3[d] | 232±1.3 | 832±1 | 259±1 | 241±1.2 | 202±1.3 | 268±1.6 | 287±1.3 | 223±1.4 | 239±1.4 | 1306±1.4 | 238±1.5 |

[a]The dilution rate of the sera that kill 50% of *B. burgdorferi* B31-A3. Shown is the mean ± standard deviation of the $BA_{50}$ values derived from three experiments (three replicates per experiment).

[b]Sera from the mice immunized with Untagged CspZ-YA

[c]Sera from the mice immunized with histidine tagged CspZ-YA

[d]1, 2, and 3 indicate the mice immunized with indicated antigen once, twice, and three times, respectively, as described in Fig. 1.

**Table S3. The thermostability of CspZ-YA proteins.**

| | CspZ-YA | | His-CspZ-YA | |
|---|---|---|---|---|
| | **-** | **C187S** | **-** | **I183Y** |
| **Tm (ºC)[a]** | 58.46±0.26 | 62.72±0.17 | 57.58±1.13 | 61.87±0.09 |

[a]Shown is the mean ± standard deviation of the Tm values derived from six experiments (one replicate per experiment).

[b]Histidine-tagged CspZ-YA

**Table S4. Strains and plasmids used in this study.**

| Strain or plasmid | Genotype or characteristic | Source |
|---|---|---|
| <u>*B. burgdorferi*</u> | | |
| B31-A3 | Clone A3 of *B. burgdorferi* B31 isolated from *I. scapularis* ticks in US. | [80] |
| <u>*E. coli*</u> | | |
| BL21(DE3) | F−, *ompT hsdSB* (rB− mB−) *gal dcm* (DE3) | Novagene |
| BL21(DE3)/pET28a-CspZ | BL21(DE3) producing residues 19 to 237 of CspZ from *B. burgdorferi* B31-A3 | [30] |
| BL21(DE3)/pET28a-CspZ-YA | BL21(DE3) producing residues 19 to 237 of CspZ with tyrosine-207 and -211 simultaneously replaced by alanine residues | [30] |
| BL21(DE3)/pET41a-CspZ-YA | BL21(DE3) producing residues 19 to 237 of CspZ with tyrosine-207 and -211 simultaneously replaced by alanine residues | [61] |
| BL21(DE3)/pET41a-CspZ-YA$_{C53S}$ | BL21(DE3) producing residues 19 to 237 of CspZ-YA with cysteine-53 replaced by serine | This study |
| BL21(DE3)/pET28a-CspZ-YA$_{T67P}$ | BL21(DE3) producing residues 19 to 237 of CspZ-YA with threonine-67 replaced by proline | This study |
| BL21(DE3)/pET28a-CspZ-YA$_{I80T}$ | BL21(DE3) producing residues 19 to 237 of CspZ-YA with isoleucine-80 replaced by threonine | This study |
| BL21(DE3)/pET28a-CspZ-YA$_{F105P}$ | BL21(DE3) producing residues 19 to 237 of CspZ-YA with phenylalanine-105 replaced by proline | This study |
| BL21(DE3)/pET28a-CspZ-YA$_{I115T}$ | BL21(DE3) producing residues 19 to 237 of CspZ-YA with isoleucine-115 replaced by threonine | This study |
| BL21(DE3)/pET28a-CspZ-YA$_{K136E}$ | BL21(DE3) producing residues 19 to 237 of CspZ-YA with lysine-136 replaced by glutamate | This study |
| BL21(DE3)/pET28a-CspZ-YA$_{V142M}$ | BL21(DE3) producing residues 19 to 237 of CspZ-YA with valine-142 replaced by methionine | This study |
| BL21(DE3)/pET28a-CspZ-YA$_{I183Y}$ | BL21(DE3) producing residues 19 to 237 of CspZ-YA with isoleucine-183 replaced by tyrosine | This study |

| | | |
|---|---|---|
| BL21(DE3)/pET41a-CspZ-$YA_{C187S}$ | BL21(DE3) producing residues 19 to 237 of CspZ-YA with cysteine-187 replaced by tyrosine | This study |
| BL21(DE3)/pET28a-CspZ-$YA_{G139M}$ | BL21(DE3) producing residues 19 to 237 of CspZ-YA with glycine-139 replaced by methionine | This study |
| <u>Plasmids</u> | | |
| pET28a-CspZ | KanR[a]; pET28a encoding protein residue 19 to 237 of CspZ from *B. burgdorferi* B31-A3 | [30] |
| pET28a-CspZ-YA | KanR; pET28a encoding protein residue 19 to 237 of CspZ with tyrosine-207 and -211 simultaneously replaced by alanine residues | [30] |
| pET41a-CspZ-YA | KanR; pET41a encoding protein residue 19 to 237 of CspZ with tyrosine-207 and -211 simultaneously replaced by alanine residues | [61] |
| pET41a-CspZ-$YA_{C53S}$ | KanR; pET41a encoding protein residue 19 to 237 of CspZ-YA with cystein-53 replaced by serine | This study |
| pET28a-CspZ-$YA_{T67P}$ | KanR; pET28a encoding protein residue 19 to 237 of CspZ-YA with threonine-67 replaced by proline | This study |
| pET28a-CspZ-$YA_{I80T}$ | KanR; pET28a encoding protein residue 19 to 237 of CspZ-YA with isoleucine-80 replaced by threonine | This study |
| pET28a-CspZ-$YA_{F105P}$ | KanR; pET28a encoding protein residue 19 to 237 of CspZ-YA with phenylalanine-105 replaced by proline | This study |
| pET28a-CspZ-$YA_{I115T}$ | KanR; pET28a encoding protein residue 19 to 237 of CspZ-YA with isoleucine-115 replaced by threonine | This study |
| pET28a-CspZ-$YA_{K136E}$ | KanR; pET28a encoding protein residue 19 to 237 of CspZ-YA with lysine-136 replaced by glutamate | This study |
| pET28a-CspZ-$YA_{V142M}$ | KanR; pET28a encoding protein residue 19 to 237 of CspZ-YA with valine-142 replaced by methionine | This study |
| pET28a-CspZ-$YA_{I183Y}$ | KanR; pET28a encoding protein residue 19 to 237 of CspZ-YA with isoleucine-183 replaced by tyrosine | This study |
| pET41a-CspZ-$YA_{C187S}$ | KanR; pET41a encoding protein residue 19 to 237 of CspZ-YA with cystein-187 replaced by serine | This study |

| | | |
|---|---|---|
| pET28a-CspZ-YA$_{G139M}$ | KanR; pET28a encoding protein residue 19 to 237 of CspZ-YA with glycine-139 replaced by methionine | This study |

[a] Kanamycin resistant
