## Supplementary figures and images for "Mechanistic insights into structure-based design of a Lyme disease subunit vaccine"

### Figure S1

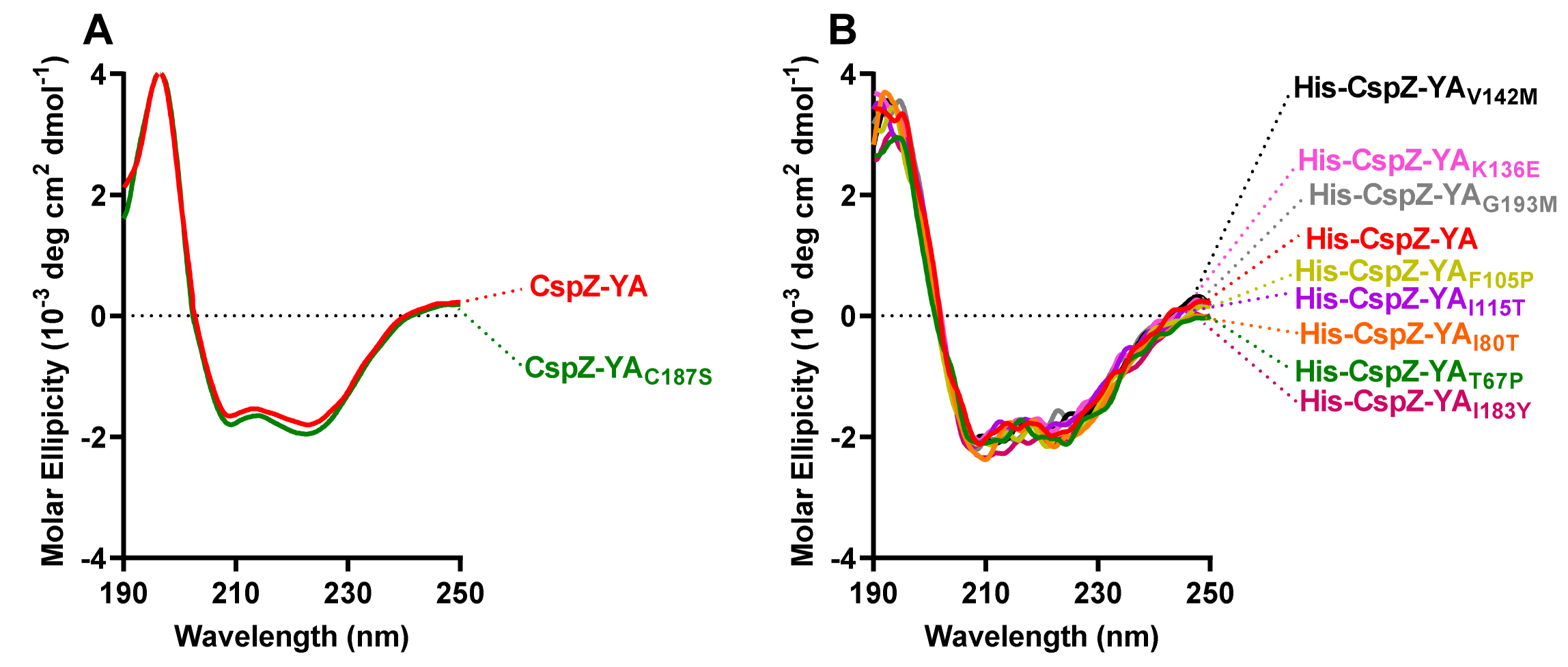

### Figure S2

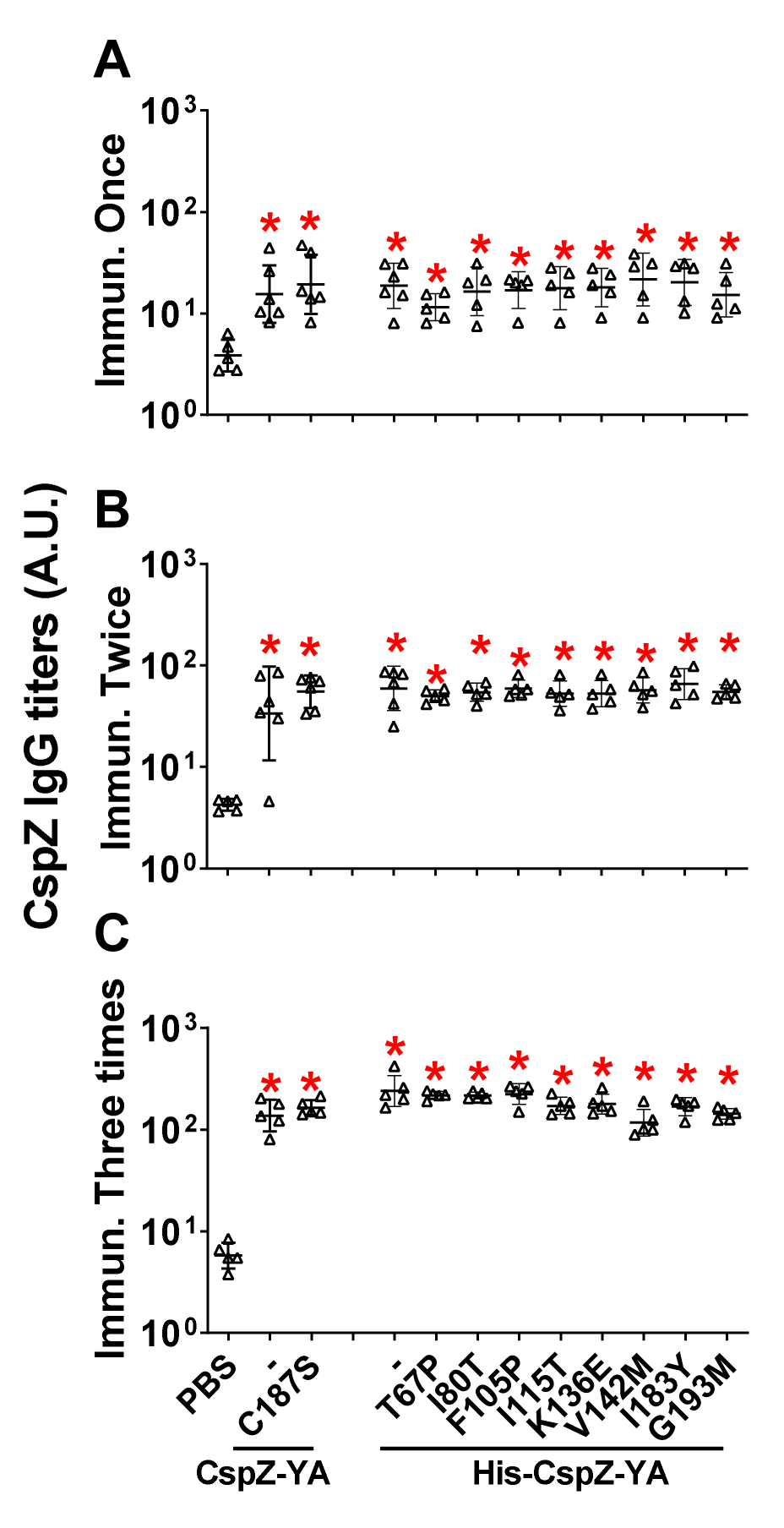

### Figure S3

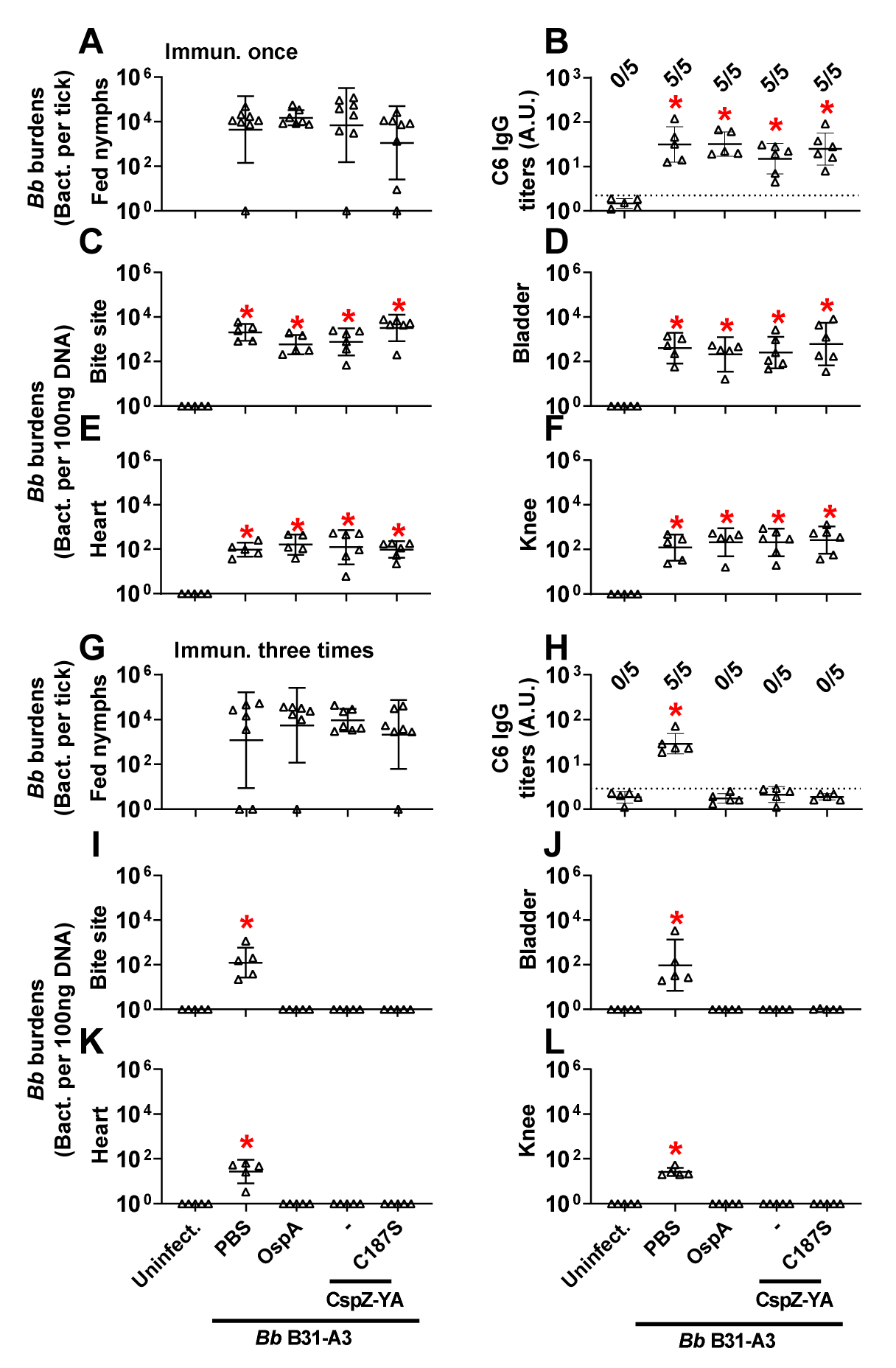

### Figure S4

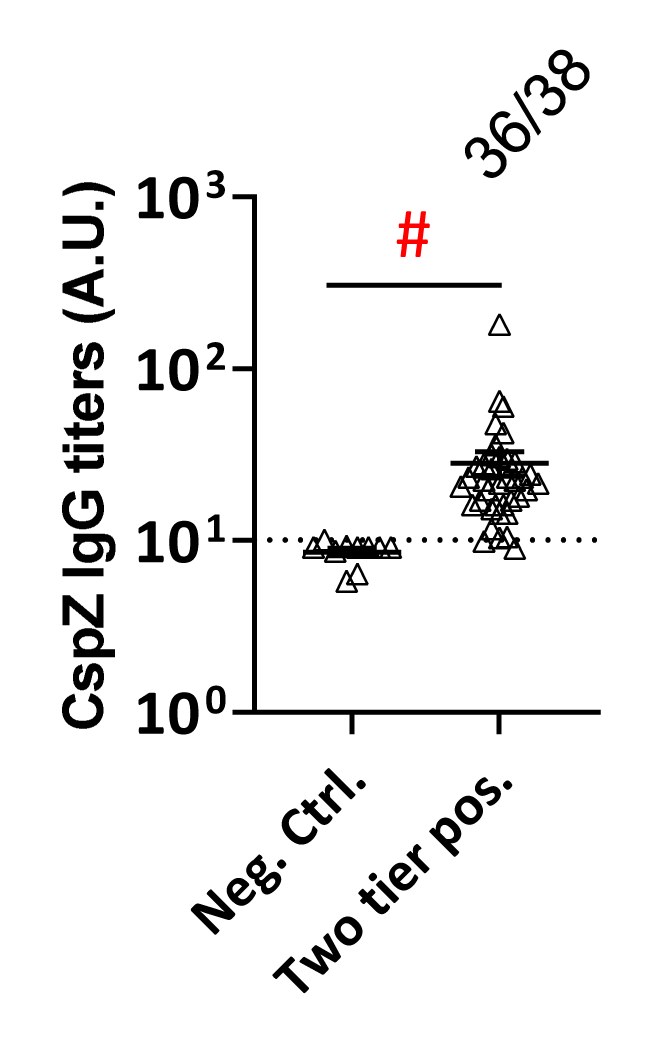

### Figure S5

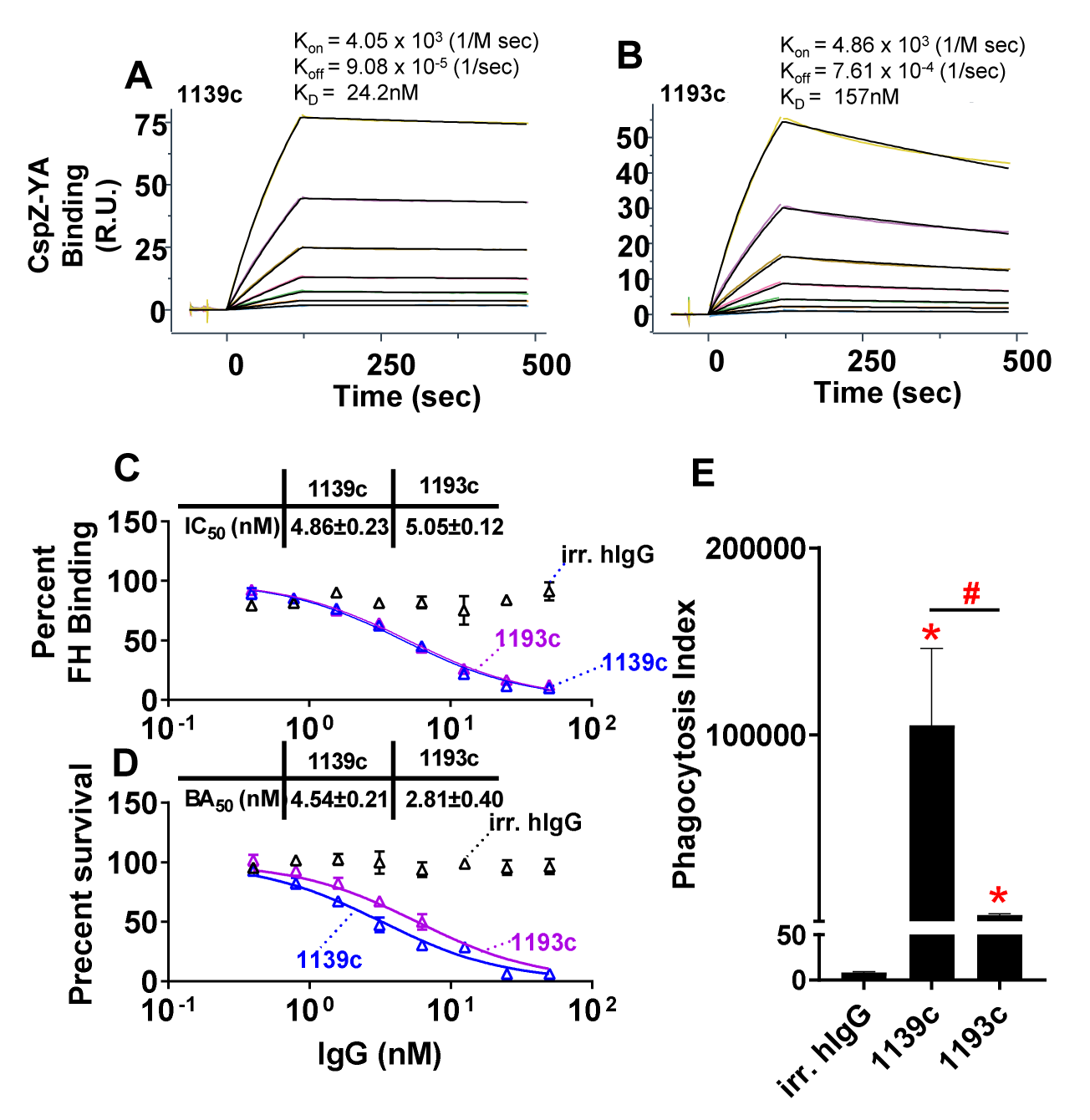

### Figure S6

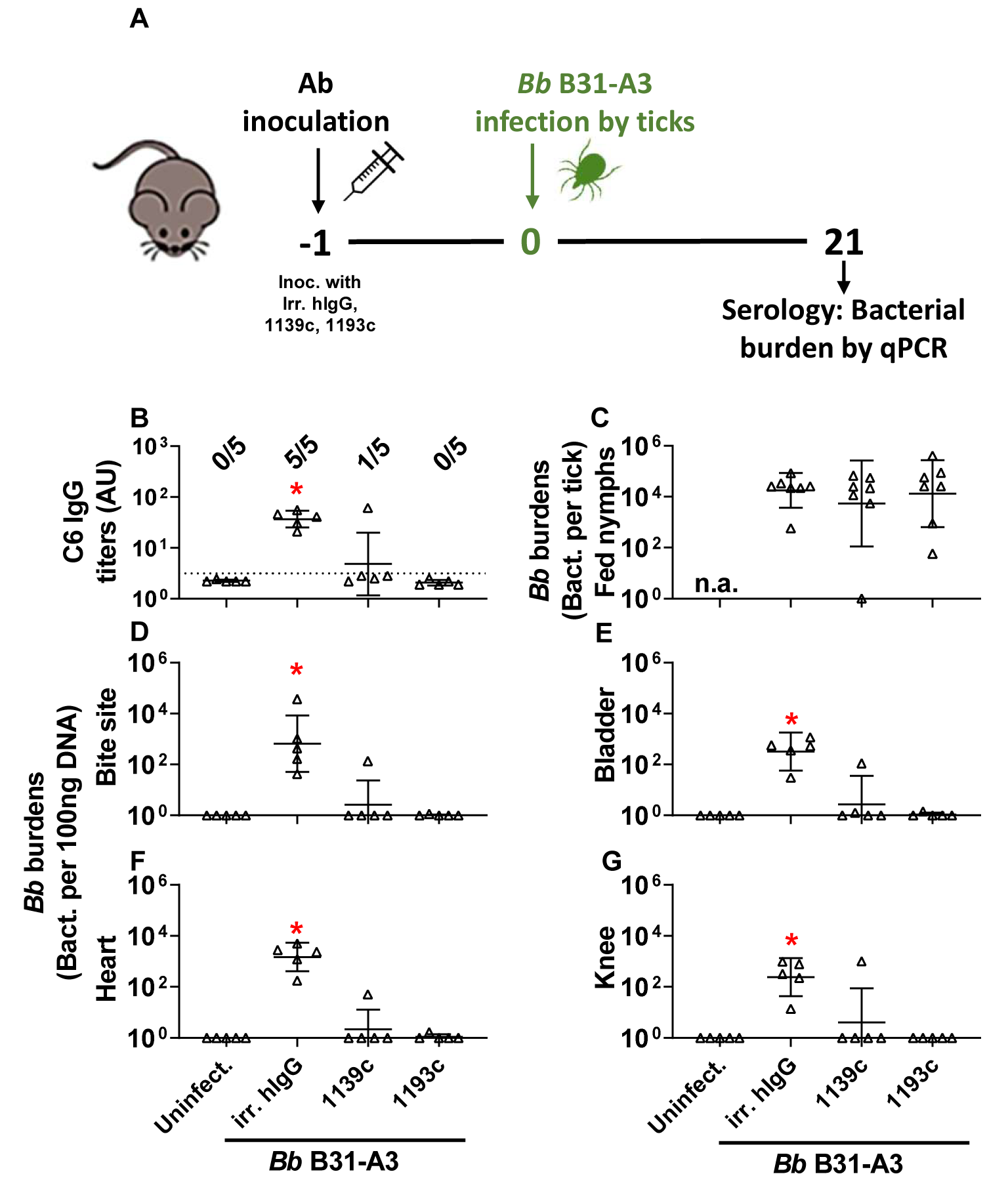
